# Age-related changes in proprioception are of limited size, outcome-dependent and task-dependent

**DOI:** 10.1101/2025.05.22.655043

**Authors:** Stien Van De Plas, Jean-Jacques Orban de Xivry

**Affiliations:** Departement of Movement Sciences, KU Leuven, Leuven, Belgium; Leuven Brain Institute, KU Leuven, Leuven, Belgium

**Keywords:** aging, proprioception, touch, recalibration, multisensory integration

## Abstract

Our ability to sense the position and movement of our limbs is essential for all activities of daily living. This ability arises from the signal sent by muscle spindles to the brain. While there is clear evidence for age-related changes in the quantity of muscle spindles and in their sensitivity, behavioral assessment of age-related changes in position sense have produced mixed findings even though it is taken as textbook knowledge that proprioception declines with age. Yet, study results are difficult to compare since there is no golden standard for assessment of proprioception. Therefore, we measured upper limb proprioception across several standard proprioceptive tasks together with key factors that could influence behavioral results such as touch, motor function, and cognition in 37 young (19-32 years old) and 35 older (53-71 years old) adults. We tested age-related differences in behavioral outcomes and their associations across tasks. Our results showed that age-related effects were variable, ranging from tasks where older participants performed better to tasks where they exhibit large age-related declines. Age-related declines in position sense are outcome-dependent, sometimes requiring large samples to detect small effects. The results were confirmed by meta-analysis based on data from hundreds of participants tested in our laboratory on the exact same tasks. Associations between outcome variables across or within proprioceptive tasks were overall negligible to weak. In conclusion, age-related changes in proprioception are limited in size, task- and outcome-dependent, and current tasks used to assess proprioception do not provide consistent evidence of age-related impairment in upper limb proprioception.

## Introduction

Throughout our daily lives we perform many goal directed movements without always looking at our body or our limbs. For example, when cooking, we pick up a pasta box and move it from one hand to the other without continuously keeping our eyes on our hands. The smoothness of this transfer of the object is possible thanks to our ability to perceive the position and motion of our limbs, called proprioception. Proprioception is intertwined with motor function, and a deficit in proprioception can have a large influence on motor actions (Allum et al., 1998; Dietz, 2002; Gentilucci et al., 1994; Gordon et al., 1995; Moon et al., 2021; Proske & Gandevia, 2012; Rossignol et al., 2006). Receptors for proprioception, also called proprioceptors, are located in the muscles, joints and tendons. Muscle spindles, which are considered the main proprioceptors for position and movement sense, are sensitive to changes in muscle length. They are located in the intrafusal fibers of our muscles (Proske & Gandevia, 2012). With age, the amount of intrafusal fibers decreases in humans (Swash & Fox, 1972), and studies in rats have found that spindles get denervated (Desaki & Nishida, 2010) or lose their typical configuration with age (Kim et al., 2007). Given the importance of proprioception to remain functionally independent with age in everyday life, we need clear insights into age-related declines in proprioceptive function.

It is considered textbook knowledge that proprioception deteriorates with age (Butler et al., 2008; Goble et al., 2009; Herter et al., 2014; Landelle et al., 2021; Lord et al., 1994; Lord & Ward, 1994; Proske & Gandevia, 2012; Ribeiro & Oliveira, 2007; Shaffer & Harrison, 2007; Tulimieri & Semrau, 2023). Some studies, however, found that proprioceptive abilities are preserved with age (Coffman et al., 2021; Djajadikarta et al., 2020; Dunn et al., 2015; Goble et al., 2012; Jordan, 1978; Pickard et al., 2003; Saenen et al., 2022; Schmidt et al., 2013; Weijs et al., 2024). There is also some evidence that a decline is only present under certain conditions such as active movement (Carranza et al., 2023), or that a decline is only found when performance is operationalized as bias, but not precision (Van de Winckel et al., 2017; Vandevoorde & Orban de Xivry, 2021). Bias refers to the signed error away from the target while precision refers to the inter-trial variability in the estimates of limb position. Moreover, due to sampling variability, age-related differences are sometimes not replicated by the same authors (Vandevoorde & Orban de Xivry, 2021). These contradictory findings across tasks, across outcomes, and across samples make it difficult to have a coherent idea about the effect of aging on proprioception. Because studies on proprioception use different tasks and measurements in different samples, the interpretation of the available evidence is complicated. To address this knowledge gap, we aim at studying age-related effects on proprioception with different tasks in the same sample.

The wide variety of tasks used in different studies to measure position and movement sense, include the use of multiple methods such as detection, reproduction and discrimination of positions, movements, trajectories, and velocities (Han et al., 2016; Hillier et al., 2015; Horváth et al., 2023). Static position sense is often measured with a matching task, in which one arm is passively moved toward a position and is actively matched with the contralateral arm or with the ipsilateral arm after remembering the target position (Dukelow et al., 2010; Goble, 2010), or with a discrimination task in which the participant reports if and how one position differs from another (Han et al., 2013; Lönn, 2001; Tan et al., 2007; Yang et al., 2020). Position matching tasks are considered as the gold standard and are most frequently used to study proprioception. Dynamic position sense is often assessed with reaching tasks, where the participant makes goal-directed movements towards external targets (Suetterlin & Sayer, 2014). Alternatively, proprioception can be studied with illusions. For example, vibration can induce the illusion of motion (Landelle et al., 2018, 2020), and skin stretch around the elbow can cause a shift in proprioception (Kuling et al., 2016). The rubber hand illusion, where a rubber hand is experienced as one’s own (occluded) hand through simultaneously administered sensory signals such as stroking the hand, has been used to investigate proprioceptive precision and the relation between proprioception, action, vision, touch and body ownership (Botvinick & Cohen, 1998; Chancel & Ehrsson, 2023; Kammers et al., 2009; Rohde et al., 2011). Finally, proprioceptive information can be accumulated over time and space to interpret shapes or movement patterns. This is then referred to as haptic perception (Bernstein et al., 1996; Gibson, 1966). While typical haptic object recognition tasks are highly dependent on touch sense (Dunn et al., 2015), a recently developed task focusses on the interpretation of proprioceptive information over time to recognize the shapes (Saenen et al., 2022, 2023). Even though all those tasks aim at measuring the construct proprioception, the obtained outcomes related to proprioceptive accuracy are often not correlated with each other (Horváth et al., 2023; Lowrey et al., 2020; Nagai et al., 2016). Also in the aging context, for example, Hoseini and colleagues (2015) found no significant correlations between three different proprioceptive task outcomes. The sample size, however, was insufficient to obtain reliable correlations. This leaves us questioning the existence of a generalizable proprioceptive ability.

While all the tasks mentioned above rely on the conscious reporting of proprioceptive information, it is also possible to measure proprioception without explicitly probing knowledge of the limb position. In daily life, we are usually not aware of where our limbs are located. Yet, by using both visual and proprioceptive information, our brain builds an estimate of the limb position (Limanowski & Blankenburg, 2016), highlighting the importance of combining different sources of sensory information. Multiple sensory inputs are integrated following Bayesian rules of optimal integration based on the reliability of senses used (Block & Bastian, 2011; Ernst & Banks, 2002; Gaveau et al., 2016; Ghahramani et al., 1997; Smeets et al., 2006; van Beers et al., 1999). In that case, the weight given to a sensory signal is inversely related to its reliability. When the information coming from two different sources differs, one can infer the weight of each source (and thus its reliability) by comparing how close the combined estimate is to each of the sources. Such mismatch between visual and touch or proprioceptive information can be artificially created in a virtual reality environment. Visual perturbations that create a mismatch with proprioception influence the estimate of hand position on a single trial basis (Rand & Heuer, 2020). Interestingly, older adults upweight visual input for integration relative to proprioceptive input, and this difference in integration is associated to a decline in proprioception (Risso et al., 2024).

Imposing such sensory discrepancy for many consecutive trials is typical in visuomotor rotation studies (Held & Hein, 1958; Morehead & Orban de Xivry, 2021). Many studies have reported a proprioceptive shift of the hand position after adaptation (Barkley et al., 2014; Clayton et al., 2014; Cressman et al., 2010; Cressman & Henriques, 2009; Mostafa et al., 2015; Ostry et al., 2010; Rand et al., 2013; Ruttle et al., 2016; ’t Hart & Henriques, 2016; Vachon et al., 2020). A recent theory hypothesized that this shift in the perceived hand position is causing the adaptation of the reach direction (Tsay et al., 2022) and is not a mere consequence of the adaptation. If so, the extent of the adaptation would be determined by how much the perceived hand position is shifted with respect to its actual position by the mismatch between the visual and proprioceptive signals. If adaptation results from the combination of visual and proprioceptive signals, the lower the reliability of the proprioceptive information, the larger the adaptation should be. If older adults do have worse proprioception, this theory could explain why older adults exhibit larger implicit adaptation (Cisneros et al., 2024; Vandevoorde & Orban de Xivry, 2019, 2021). While we have failed to find such a link in the past (Vandevoorde & Orban de Xivry, 2021), the question is still open.

To investigate the cause of this inter-study heterogeneity and to increase our understanding of the influence of aging on proprioception, we obtained data from five proprioceptive tasks within the same sample of 37 young and 35 older adults. These proprioceptive tasks like any others are not only influenced by the quality of the proprioceptive signaling but also by the accuracy of motor behavior, the interaction with signals coming from other senses such as touch (Delhaye et al., 2018; Kuling et al., 2016; Proske & Gandevia, 2012; Zopf et al., 2011), and by cognitive abilities (Ager et al., 2024; Oh-Park et al., 2013). To control the possible influence of these factors on our results, the participants also performed a touch discrimination task, two motor tasks and a cognitive task. Together, these data will allow us to shed light on the effect of aging on proprioception.

## Methods

### Participants

Seventy-three people between 19 and 32, or between 53 and 71 years old volunteered to participate in this study. Data of 72 volunteers were included in the analysis. They did not show any cognitive impairment as measured with the Montreal Cognitive Assessment (score ≥ 26), were right-handed as assessed with the Oldfield handedness questionnaire and had no history of neurological disease nor addiction or excessive use of caffeine/alcohol. Data of one volunteer was excluded from the analysis because the volunteer had a shoulder implant. Of the 72 included participants, 37 were young adults (age 19 to 32 years; average age 24.89 ± 3.60 years; 22 females) and 35 were older adults (age 53 to 71; average age 64.17 ± 4.49; 18 females). All volunteers signed the informed consent form. A renumeration of 20 Euro was given for participating in this study. This study was approved by the medical ethical commission of UZ/KU Leuven (study number S65248).

### Experimental setup

Participants completed two sessions of maximally two hours. In one session, data was collected with the Kinarm end-point Lab robot (BKIN Technologies Ltd., Kingston Canada). This robot had a left and right handle which the participant could manipulate in the horizontal plane, using their left and right hands. Participants’ vision of the arms and hands was blocked by a bib to prevent direct visual feedback. Instead, visual feedback about the task and the position of the hands could be provided by the virtual reality display component of the End-point Lab. This visual information could be seen on a mirror mounted horizontally above the upper limbs in front of the participant. The mirror was positioned in parallel to the movement plane and reflected the image from a screen on top of it. Eight tasks were completed in a randomized order. The ninth task was a motor adaptation task and was always completed at the end of the session to prevent interference of motor learning on performance in other tasks. In the other session, videos of upper limb movements were made with a GoPro HERO8 camera. Behavioral data of two tasks (a tactile sensation task called Bumps and a motor dexterity task called Purdue pegboard) from the video session were included in the analysis of this paper. The video data will not be discussed in this paper. The order of the two sessions was randomized between participants, which means that the Montreal Cognitive Assessment (MoCA) was sometimes done in the Kinarm session and sometimes in the video session.

### Assessments

#### Sensory function

The goal of this study was to find evidence for age-related differences in sensory function and how those differences are related to differences in motor function and cognition. The investigated sensory functions were proprioception and touch. Several aspects of proprioception were measured (i.e. matching, discrimination, reproduction).

##### Arm position matching

The arm position matching task is a Kinarm Standard Test available for the End-Point Lab (Dukelow et al., 2010). It assesses the position sense of the participants in the upper limbs. Participants mirror-matched the position of one hand across the body midline with their other hand (Fig.1). The arm to be assessed was passively moved by the Kinarm robot to one of four possible target positions which are spaced 20 cm apart on the corners of an invisible square (Fig.1A). The participant actively mirror-matched this position with the other hand (Fig.1B). There was no time limit on mirror-matching. To move from one trial to the next participants informed the experimenter orally when they thought their hand was in the correct position and the experimenter started the next trial. Visual feedback on hand position was not provided during this task so the participant had to rely on the felt position of the hand to complete the task. Each of the four targets was repeated 6 times, summing up to 24 trials per assessed hand and 48 trials per participant because we assessed both hands.

**Figure 1.**
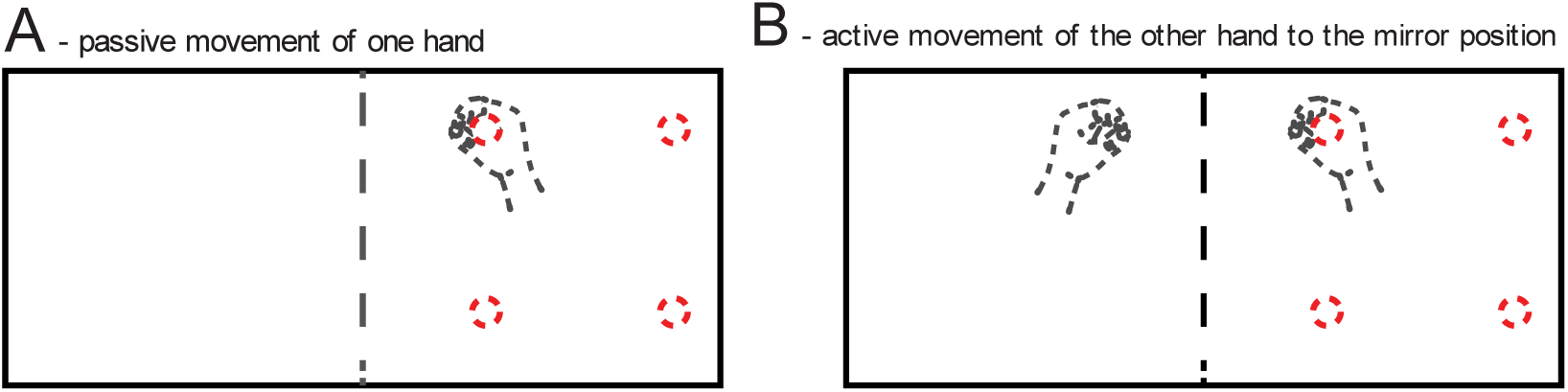
Arm position matching task. Invisible target locations are depicted as dashed red circles. A) The arm to be assessed (e.g. right arm) is passively moved by the robot to one of four targets. B) The participant actively mirror-matches the position with his/her other arm.

To quantify proprioceptive accuracy of mirror-matching in the arm position matching task, we measured the difference between hand and target position. First, the absolute error was calculated in the X and Y direction separately for each trial. Next, the mean of these absolute distance errors across the four targets was calculated as the absolute position error. The higher the absolute error, the worse the position matching accuracy.

Next to the absolute position error, we also assessed the consistency of matches across trials. This was quantified by the inter-trial variability of the matched positions. First, the mean standard deviation in the X and Y direction were calculated for all targets. Next, the root-mean-square of these mean standard deviations in the X and Y direction was defined as the inter-trial variability. The higher the inter-trial variability, the higher the spread of mirror-matches around a target, the less consistent a participant was in judging the position of the hand from one trial to the next.

##### Perceptual boundary

The perceptual boundary task was used by Vandevoorde and Orban de Xivry (2021) who made an adapted version of the task by Ostry et al. (2010). In this task (Fig.2) participants used their right hand to operate the Kinarm robotic manipulandum. The participants were first instructed to reach from the center of a circle to a target straight ahead. The target was represented as an aperture in the circle. While the participant was actively reaching from the starting point (red square) to the target without any visual feedback about the hand position, the robot constrained the movement of the hand to a straight line that was tilted to the left or to the right by a certain angle (dashed blue rectangle in Fig.2B). The robot’s action pushed the hand away from the target, and the participants were asked to report verbally whether their hand was at the left or right of the target.

**Figure 2.**
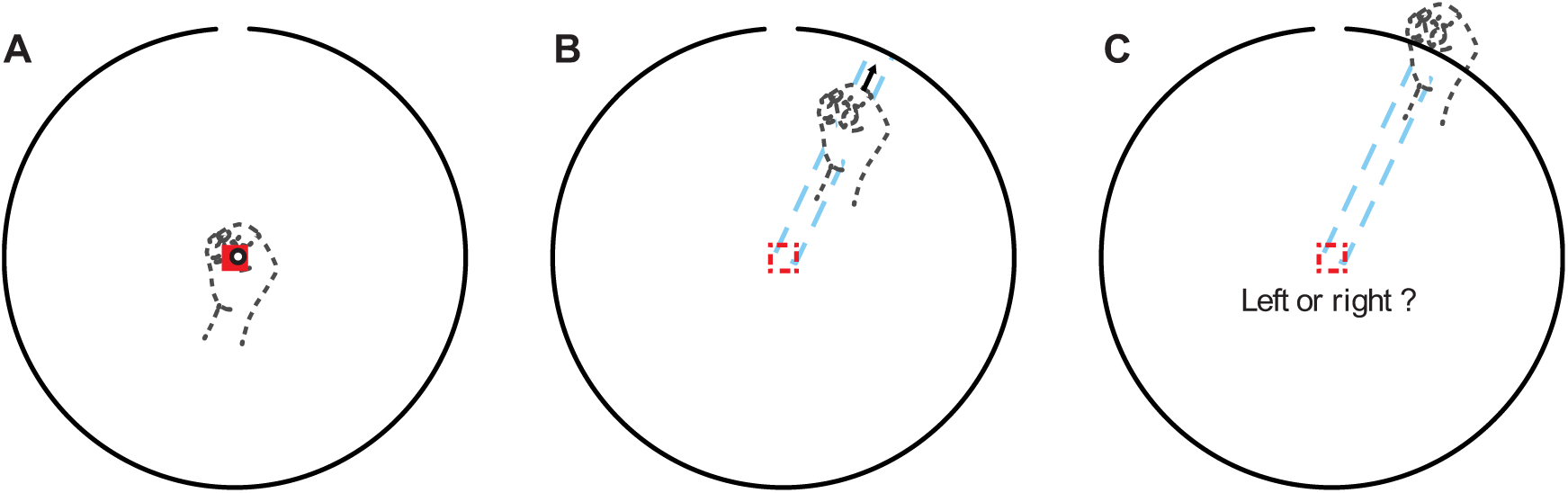
Perceptual boundary task. A) The participant moves the visible cursor (white dot) representing the hand position into the starting point (red square) to make the target appear (circle with aperture). B) The cursor and starting target are no longer visible (dashed circle and dashed square respectively). During movement, the hand is constrained by a channel (dashed blue rectangle) to deviate the hand from the target by a given angle. C) At the end of the trial, participants verbally report to which side they think their hand was displaced (left or right of the target).

Performance accuracy was estimated by a parameter by sequential testing (PEST) procedure (Taylor and Creelman, 1967). This procedure started with a large deviation (25° or 23.5° to the left or to the right) relative to the straight line from start to target (defined as the 0° line). The direction and the magnitude of the deviation were changed with each trial depending on the answer of the participant. The angle of the deviation was changed in the direction opposite to the answer of the participant. If s/he said left, the angle was changed towards the right by an angle defined by the step size. The step size started at 16° and was divided by 2 each time the participant changed their answer (answer “left” in trial t and “right” in trial t+1, or the other way around). Once the step size fell below a minimum threshold of 1.5°, the procedure was ended. Each participant completed 8 PEST procedures, two per deviation magnitude (25 or 23.5°) and direction (left or right). The number of movements per block differed per participant due to the adaptive nature of the procedure.

The data of all 8 PEST procedures were used to fit a psychometric curve from which two parameters were derived to quantify proprioceptive accuracy. The psychometric curve was calculated as follows. First, a generalized linear regression model was fitted with function “glmfit” in Matlab. Because this task generated binary responses (answer “left” or “right” for each trial), we fitted a sigmoidal probability response curve for binomially distributed data, with the ‘logit’ link function:

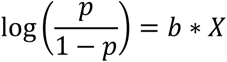

The Matlab function returned the coefficient estimates, a 2-by-1 vector *b*, which were based on the response variables, *p*, and the predictor variables, *X*. *p* was defined as a 101-by-2 matrix with the number of successes in the first column and the number of trials in the second column for each of the possible angular displacements. *X* was a 101-by-2 matrix including a constant term in the first column (column of ones) and the displacement angles in the second column (−25° to 25° in steps of 0.5°). Second, the coefficient estimates, b, where used with the function “glmval” to predict the values of the psychometric curve for any angular displacement between −25° and 25° that had not been tested.

From the psychometric curve we calculated two parameters. The proprioceptive sensitivity was calculated as the slope of the straight line between the 25% and 75% probability and represented the ability to discriminate between angular displacements during reaching. The higher the slope, the more sensitive a person is to small differences in perturbation angles. Second, the bias represented which direction subjectively felt like 0° displacement from target, i.e. which direction felt like reaching straight ahead. This bias to the left/right was calculated as the displacement angle (x-value) corresponding to the inflection point of the psychometric curve (50% probability of answering “right”). A positive bias indicates a bias towards the right, a negative one to the left.

##### Shape reproduction

The passive version of the sensory processing task developed by Saenen et al. (2022) was used. The task includes passive exploration, active reproduction and visual identification of geometrical shapes (Fig.3). During exploration, the Kinarm moved the dominant (right) arm passively so that it traced one of the shapes. Subsequently, the participant was instructed to actively reproduce that shape with the non-dominant (left) arm with the same speed and the same size. At the end of each trial, the participant was shown six different shapes out of which s/he indicated the shape just traced during exploration and reproduction (identification phase). Each participant completed one practice trial to get familiar with the task, followed by 16 test trials.

**Figure 3.**
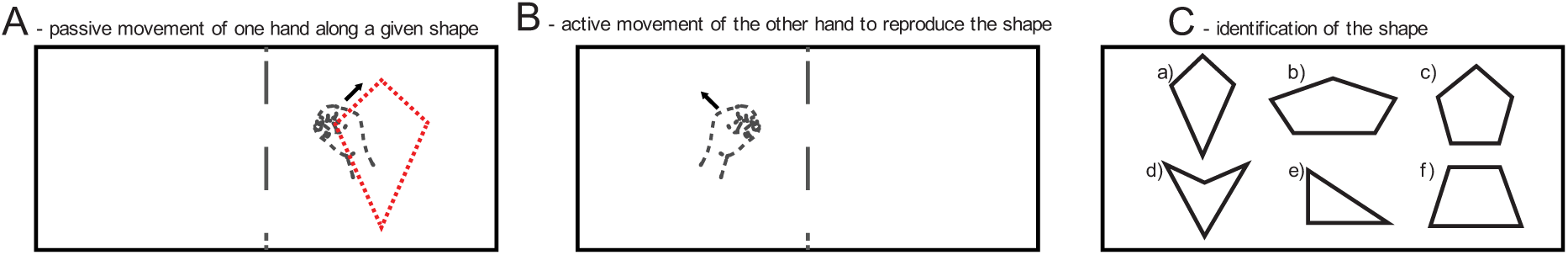
Shape reproduction task. A) Exploration phase: the participant brings the right hand to the starting point and the hand is passively moved by the robot in a given shape. The invisible cursor is illustrated by a dashed circle. B) Reproduction phase: the participant brings the left hand to the starting point and actively reproduces the shape. The visible cursor is illustrated by a circle. C) Identification phase: the participant verbally identifies the shape out of six options.

In contrast to the study by Saenen et al. (2022), participants were instructed to mirror the shapes during the reproduction phase. All shapes are symmetric, but the direction of movement is different between arms. For parameter calculation, mirrored trials were mirrored back again to ensure optimal shape overlap.

In the shape reproduction task, proprioceptive accuracy was quantified by means of four parameters on which an exploratory factor analysis with one factor was performed, as described in Saenen et al. (2022). The first two parameters quantified the similarity between the explored (passive movement imposed by the robot) and the reproduced (active movement by the participant) shapes by means of cross-correlation for each direction separately. To do so, the data from the exploration and reproduction phases were first normalized in time by means of resampling via linear interpolation (35 000 samples) for each phase. Then, the maximal cross-correlation between the two signals was computed. The bigger the similarity between the two signals, the higher the value of the cross-correlation. Relative differences in timing between active reproduction compared to passive exploration resulted in lower values for cross-correlation. The cross-correlation parameter had a value between −1 and 1 with higher positive values meaning better shape reproduction.

The third parameter also quantified similarity but only considered spatial information, not time. This parameter for shape reproduction accuracy was calculated by means of Procrustes analysis. After finding an optimal superimposition between the explored and the reproduced shape by means of translating, scaling, reflecting and rotating the shape, the distance between the superimposed shapes was calculated as the square root of the sum of the squared distances between the corresponding points of the two shapes. Lower values of the parameter reflected less distance between the shapes and thus better shape reproduction. The fourth parameter was the percentage of correctly identified shapes in the identification phase. An overall score for shape reproduction was calculated by means of an exploratory factor analysis on all four parameters (see ‘statistical analysis’ section for details).

##### Bumps

The Bumps test was developed and validated by Kennedy and colleagues (2011) to quantify tactile sensation in the finger pads. Five disks each contained, five possible target locations that consisted of 2×2cm glass plates. In the middle of one of the five plates, a cylinder with a diameter of 0.5 mm and a height of 5, 10, 15, 20 or 25 µm could be found. The participant had to identify which of the five glass plates contained the small bump. Participants completed three repetitions for each bump height, summing up to 15 trials per participant.

Tactile sensation in the finger pad of the right index finger was quantified as a bump threshold. The threshold was calculated as the bump height for which the participant can correctly detect the bump in 2 out of 3 repetitions (following Weber’s law of absolute threshold).

#### Integration/recalibration with vision

While position sense can be reported consciously by participants, it can also be used by the motor system unconsciously. In such a case, the influence of proprioception can be measured indirectly via movement kinematics, which are influenced by both visual and proprioceptive feedback. If a mismatch between the two senses occurs, a weighted average of both the state estimate from vision and proprioception is performed following Bayes rules. In this case, the uncertainty of each of the senses determines its relative weight into the estimate.

##### Task-irrelevant clamped feedback

The task-irrelevant clamped feedback task is an implicit motor adaptation task where the extent of motor adaptation has been postulated to be dependent on the quality of the proprioceptive information about the estimated movement direction (PReMo model, Tsay et al., 2022) even though some studies casted doubt on this link (Tsay et al., 2024; Vandevoorde & Orban de Xivry, 2021).

In the task-irrelevant clamped feedback task, the participant reached from a central target towards one of four peripheral targets placed at a distance of 8cm from the central target (Fig.4). S/he was instructed to move through the target in a straight line and return to the center. A point was given each trial the target was reached between 200 and 300ms after onset of movement. Visual feedback was given by means of a round white cursor on the Kinarm visual display. At the point where the hand crossed the target, the cursor took on a square form and remained stationary for 1.5 seconds to give feedback on reaching accuracy. In the first 20 trials, defined as the baseline phase, the cursor motion reflected the actual hand motion. In a practice phase, participants could get familiar with the concept of clamped feedback. Clamped feedback means that the direction of cursor motion is fixed at an angle relative to the straight line from start to target (dashed line in Fig.4B). The cursor will move independent of hand direction, but the velocity of the cursor remains matched to the velocity of the hand. Participants performed 6 reaches towards a target with visual feedback clamped at 3.5°. They practiced ignoring the cursor by reaching in different directions without seeing any change in the direction of cursor motion. The data of the practice phase were not included in the analysis. Afterwards, visual feedback was clamped at 30° to the left of the target. Participants were instructed to ignore the cursor motion and to continue bringing their invisible hand to the target. They did so for 240 trials with a one-minute break every 80 trials.

**Figure 4.**
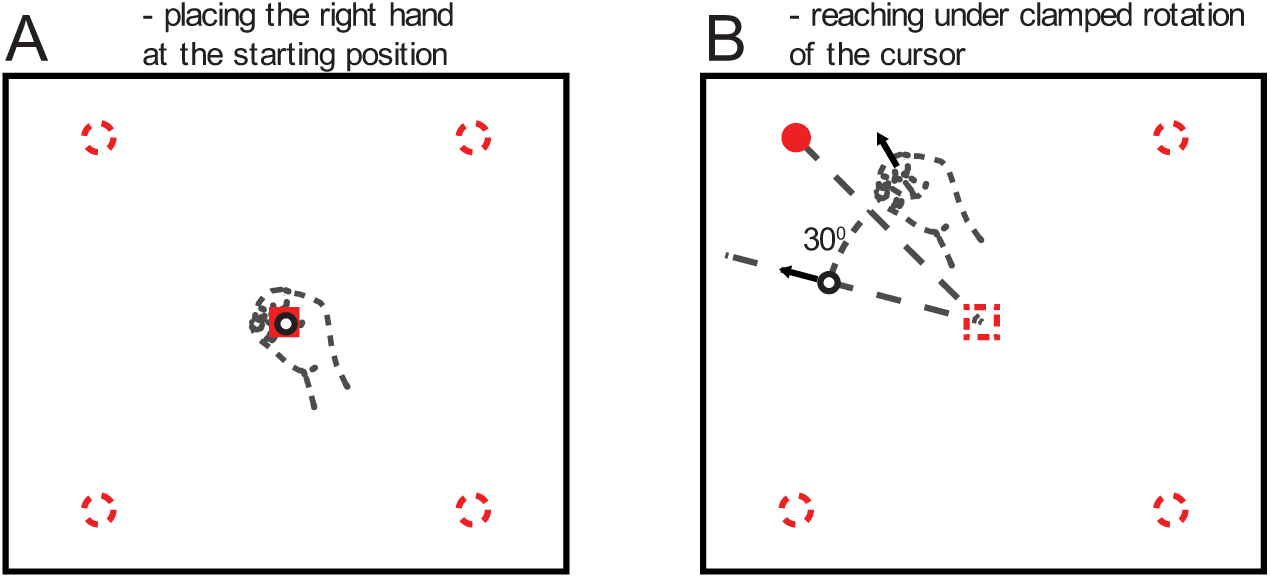
Task-irrelevant clamped feedback task. A) The participant moves the hand into the central starting position (red square). The hand location is visible as a cursor (black and white circle). Peripheral targets (dashed red circles) are not yet visible. B) One peripheral target becomes visible (red dot). The starting point (dashed red square) and actual hand location (dashed black circle) become invisible. The cursor motion (trajectory of the cursor is represented by the dashed black line under the black and white circle representing the cursor) always goes 30° to the left with respect to the trajectory straight to the target (dashed black line), independently of the actual hand trajectory. This visible but clamped cursor is represented by black circle.

Clamped feedback gives rise to implicit adaptation (Morehead et al., 2017). We quantified the level of implicit adaptation as the mean hand angle during the last 30 trials of the test phase, corrected for baseline. The hand angle in one trial was defined as the angle between the straight line from the central target to the peripheral target and the straight line from the central target to the position of the hand at 4 cm distance from the central target. The mean hand angle of baseline was subtracted from the hand angles of the test trials. Bigger hand angles indicated higher levels of implicit adaptation to clamped visual feedback.

##### Integration of proprioception with vision

The integration of proprioception with vision task is a reaching task based on the work by Rand and Heuer (2020) and aims at measuring the relative contribution of visual and proprioceptive information on arm position sense (Fig.5). The main idea behind the task is that participants have to use an internal estimate of the hand position in order to move to a given target. This internal estimate theoretically arises from a weighted average of a visual estimate and a proprioceptive estimate. The weights of these averages are inversely related to the reliability of the sensory signals.

**Figure 5.**
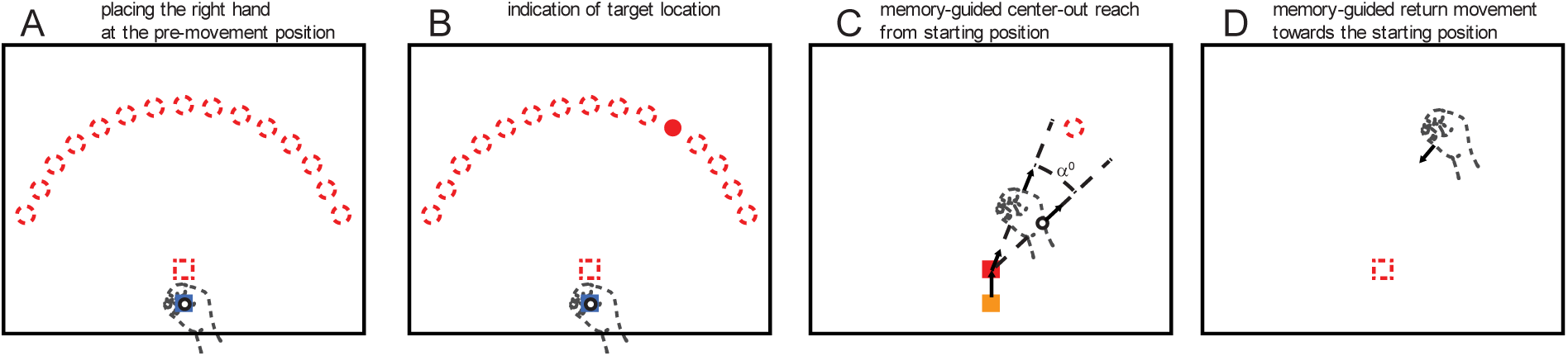
Integration of proprioception and vision task. A) The pre-movement target appears (blue square). The 15 possible target locations are represented as dashed red circles. Participants are instructed to bring the visible hand cursor into the pre-movement target. B) Moving the cursor into the pre-movement target prompts the peripheral target (red dot) to appear. The target remains visible for one second. C) The target disappears from the screen (red dashed circle) while the starting point (red square) appears, and the pre-movement target turns orange as a cue to move into the starting target. As soon as it is reached, the starting target disappears, and the participant continue to move towards the remembered location of the target (black arrows). In 25% of the trials, the hand cursor position (solid black circle) is rotated from the actual hand location (dashed hand). The rotation angle α was −25°, −15°, +15°, or +25°. In 75% of the trials, the hand cursor position matched the actual hand position (not shown, α = 0°). D) After reaching to the remembered target, the participant returns towards the remembered starting target position (red dashed square) in the absence of any new visual information about the hand location at the end of the reach or about the position of the starting target.

In this task, each trial starts with the appearance of a pre-movement target (blue square in Fig.5A-B) and the participant was instructed to move the cursor, a white dot, into this pre-movement target. This action prompted the peripheral target (red dot in Fig.5B) to appear, and the participants were instructed to memorize this target position. There were 15 possible target positions (dashed red circles in Fig.5A), all located 15 cm from the starting target. After 1 second, the target disappeared, and the starting point of the reaching movement appeared (red square in Fig.5C) at 3 cm above the pre-movement target. The pre-movement target turned orange as a cue for the participant to move to the starting point, which disappeared as soon as it was reached. Participants then reached to the remembered target position (Fig.5C). The hand cursor was visible throughout this center-out movement. As soon as the target was reached, the hand cursor disappeared and the hand was stopped by a physical barrier simulated by the robot, as if the hand was stopped by a pillow such that the hand stayed at the final reach position. Finally, without further information about the current hand position, participants had to return to the starting point (Fig.5D).

In 25% of the trials, the visual feedback during the center-out movement (from the starting position to the remembered target, Fig.5C) was rotated by an angle of +/-25° or +/-15°. Rotation was randomly applied in a limited number of trials to avoid adaptation. Each participant completed 208 trials. In 52 trials the visual feedback was rotated (13 trials per rotation condition).

This task was different from the task-irrelevant clamped feedback task on two important aspects. First, this task used single trial perturbations instead of perturbations in each trial. This is done to avoid continuous adaptation and to look at concurrent processing and integration of proprioceptive and visual information. Second, this task looked at the error during the return movement within the same trial instead of looking at the error in the next trial.

The main parameter of interest was the estimated hand position after reaching out toward the target. In trials with rotated feedback, we assumed that the estimated hand position was somewhere in between the actual position of the hand (based on proprioceptive information) and the final position of the cursor (based on vision) (Colonius & Diederich, 2020; van Beers et al., 1999). If more weight was given to proprioceptive information relative to visual information to estimate the hand location, then the estimated hand position should be closer to the actual hand location. If, on the other hand, more weight was given to visual information relative to proprioceptive information, the estimated hand position should be closer to the cursor location. The direction of the return movement is determined by where the motor system estimated the hand position. Therefore, we looked at how much the return movement direction differed from the expected return movement direction (defined by a straight line between the actual hand position at the end of the center-out movement and the actual starting target position).

To quantify the relative weight of vision and proprioception on the estimated hand position, the parameter was calculated as follows. First, we realigned each trial with respect to the direction of the target used in the specific trial (one of the 15 dashed red circles in Fig.5A). We define the parallel direction as the direction from the starting point towards the target and the perpendicular direction as orthogonal to this direction. In this case, positive parallel values are towards the target and negative parallel values are towards the starting point. Second, the location of the hand at the start of the return movement (timepoint t0) was defined as the point where the parallel velocity of the hand reached 20% of its minimum (maximum in absolute value). Then, the point where velocity reached its minimum (i.e. the maximum in absolute value) was defined as timepoint t1. We computed the initial direction error as the angle between the line connecting the hand position from t0 to t1 and the expected return movement direction (line from hand position at t0 to the starting target position). Finally, for each participant, the mean initial direction error was calculated for each condition (−25°, −15°, 0°, 15° and 25° rotation) and submitted to a robust linear regression with the amount of rotation as independent variable. The slope of this regression fit was used as an estimate of the relative weight of vision and proprioception on hand position estimation. The steeper the negative slope, the larger the weight given to visual information or the smaller the weight given to proprioceptive information to estimate the position of the hand.

#### Motor function

##### Visually guided reaching

The visually guided reaching task is a Kinarm Standard Test available for the End-Point Lab that aims at measuring visuomotor capabilities of the upper limb (Coderre et al., 2010) (Fig.6). The task started once the hand was in the central starting point. As soon as one of four targets became visible, the participant reached out to it as quickly and as accurately as possible and stopped inside the target. After each reach out movement, the starting point became visible again, and participants moved back to it. The distance from the central starting point to each target was 10cm. Targets were spaced 90° apart. Participants reached out five times to each of the four targets, which added up to a total of 20 reach-out movements. Only data of the reach-out movement were used for analysis.

**Figure 6.**
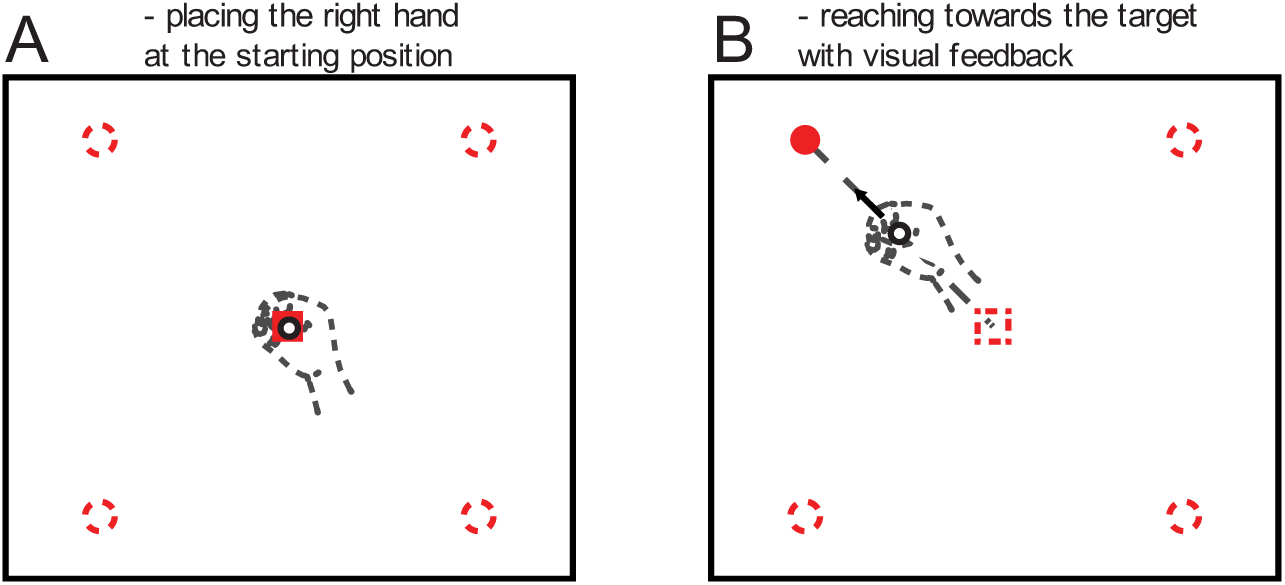
Visually guided reaching task. A) The participant brings the hand (represented by a black circle) towards the central starting point (red square) to start the trial. B) One of four possible targets (red dashed circles) becomes visible (red dot). The participant moves as quickly and accurately as possible to stop inside the target.

To quantify whole-arm visuomotor abilities, we used the parameters provided by the Dexterit-E 3.9 software of the Kinarm. We performed an exploratory factor analysis on seven of the ten parameters of the visually guided reaching task as was done by Saenen et al. (2023). The two extracted factors represented motor control and speed.

##### Purdue pegboard

The Purdue pegboard task aims at assessing fine motor skills of the fingers and gross motor skills of the upper limb. It was developed by Tiffin & Asher (1948). Participants used their dominant (right) hand to place as many pins as possible into the holes of the wooden board for 30 seconds. Pins were in a cup at the top of the board. On the board, there were two parallel columns of 25 holes. Participants were instructed to fill the right row with pins by starting at the top and not skipping any holes. Motor skills were quantified by the number of pins placed correctly onto the Purdue pegboard.

#### Cognition

##### Spatial working memory

The visual-spatial working memory task aims at quantifying working memory capacity (Christou et al., 2016; McNab & Klingberg, 2008; Vandevoorde & Orban de Xivry, 2020) (Fig.7). A circular array (diameter 6.25 cm) of 16 squares (1.9 cm x 1.9 cm) was shown on the Kinarm visual display. In three to six of the squares red dots (targets) were presented for two seconds followed by a three second interval during which the participant was instructed to focus on the central fixation cross. The circular array of squares then re-appeared with a question mark in one of the squares together with a “yes” and “no” box. Participants were asked if this square had contained a target before and had three seconds to answer the question. Participants were warned that the “yes” was sometimes placed on the left and sometimes on the right. A total of 48 trials were randomly presented but evenly distributed over the conditions with three, four, five or six targets (12 trials/condition).

**Figure 7.**
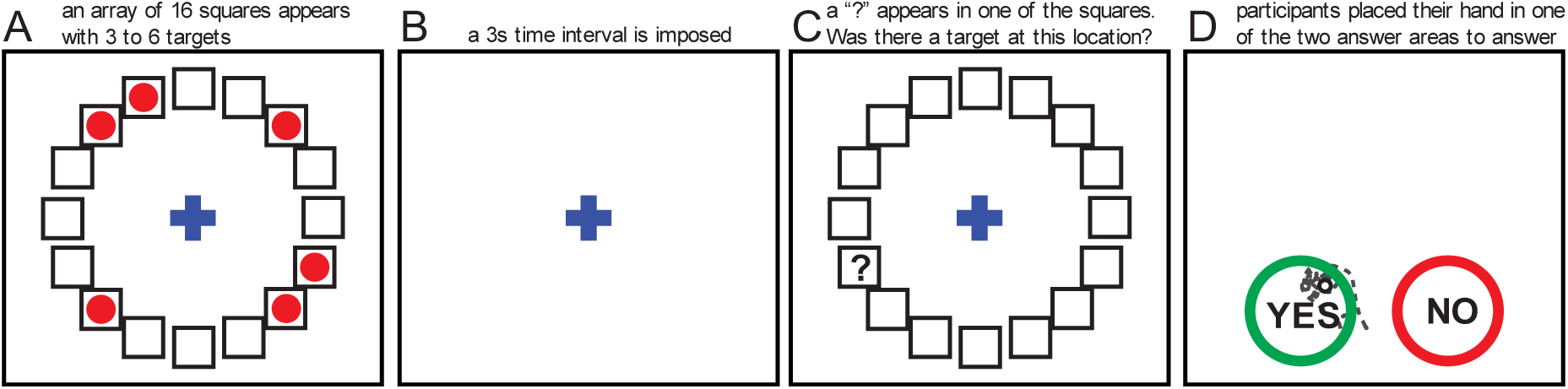
Visual-spatial working memory task. A) Targets (red dots) are presented in a circular array of squares. The participant is instructed to remember the target locations. B) The participant focuses on the fixation cross while remembering the target locations. C) A question mark is presented in or adjacent to one of the target positions. D) The participant answers whether there was previously a target presented at the position of the question mark by reaching to the “yes” or “no” box. The answer targets switch places between trials.

To quantify working memory capacity the number of items that can be stored was estimated by parameter K (Vandevoorde & Orban de Xivry, 2020; Vogel et al., 2005). Under the assumption that performance was correct in K out of S trials if the participant can hold K items out of an array of S items in memory, working memory capacity K was calculated by the formula *K = S*(H-F)* where *S* is the size of the array (the number of targets presented), *H* is the hit rate defined as the number of correct answers divided by the number of trials per condition, and *F* is the false alarm rate defined as the number of incorrect answers divided by the number of trials per condition. A correction for guessing was applied in this formula by subtracting the false alarm rate from the hit rate so that correct answers due to guessing were accounted for. The parameter K was calculated for each condition separately (three, four, five and six targets; K3, K4, K5 and K6 respectively). Ideally, K got bigger with an increasing number of targets (K6 ≥ K5 ≥ K4 ≥ K3) and the resulting parameter K was equal to the value of K6. As long as K-values increased gradually with array size, the highest K-value was chosen as the working memory capacity K (e.g. K = K5 if K5 ≥ K4 ≥ K3). If K-values did not increase gradually with array size (e.g. K5 ≥ K4 & K5 ≥ K3 but K3 ≥ K4), the mean of the highest K-value and the K’s of smaller array sizes was taken as the working memory capacity (e.g. K = mean(K5, K4, K3)). Such a decision tree to decide on the resulting K was used in previous studies (Vandevoorde and Orban de Xivry 2020, Saenen et al. 2022, 2023). The higher the value of K, the bigger the working memory capacity.

### Data analysis

#### Preprocessing

Kinarm data was imported into MATLAB 2021b (The MathWorks) using the freely available Kinarm Analysis Scripts. Calculation of parameters was also performed in MATLAB 2021b. A description of the data analysis for each task is provided in the previous section. After calculation, parameter values were imported in R for statistical analysis.

#### Statistical analysis

Robust statistical tests were performed in R version 4.4.0 using RStudio (Posit team, 2024). We performed robust t-tests to test for age-related differences on 20% trimmed means using the “yuen” function from the R-package “WRS2” (Mair et al., 2024; Mair & Wilcox, 2020; Yuen, 1974). An exception was made for parameters where an extra within-subjects factor was available, then we used the function “bwtrim” for robust two-way mixed ANOVA’s. This was the case for the absolute position error and the inter-trial variability of the arm position matching task with age as a between-subjects factor (young vs. older) and hand as the within-subjects factor (left vs. right), and for the cross-correlation of the shape reproduction task with age again as between-subjects factor (young vs. older) and axis-direction as within-subjects factor (cross-correlation in X direction vs. cross-correlation in Y direction). We used a statistical significance level of p < 0.05. The reported effect sizes and their 95% confidence interval were a robust alternative to Cohen’s delta calculated with the “akp.effect” function with the nboot argument set to 600 (Algina et al., 2005). To facilitate the interpretation of the Cohen’s d effect size, we report the superiority index (Common Language Effect Size, McGraw & Wong, 1992). The superiority index provides the probability that a randomly drawn datapoint from the values for the younger group is larger/better than one drawn from the values of the older group. In the absence of difference (Cohen’s d=0), the superiority index will be 50%.

The reported means were 20% trimmed means calculated using the function “mean” with the trim argument set to 0.2. Trimmed means were reported ± trimmed mean standard error calculated using the “trimse” function.

Exploratory factor analysis was performed in two tasks. We used the “fa” function from the “psych” package in R, using the principal factor solution with an oblimin transformation. For the shape reproduction task, parameters were combined into one factor representing sensory processing ability as defined by Saenen et al. (2022). For the visually guided reaching task, parameters were combined into two factors representing motor control and speed as defined by Saenen et al. (2023).

Robust multilevel correlations between variables were calculated using the “correlation” package in R (Makowski et al., 2020).

## Results

In our study, we investigated a group of young and a group of older adults across a range of somatosensory, motor and cognitive functions. Our goal was to test whether there was a consistent age-related decrease in sensory function across modality (proprioception vs. touch), outcomes (accuracy vs. consistency), and tasks. In addition, we wanted to investigate how age-related changes in somatosensory function were related to changes in the motor or cognitive domains.

### Aging has a limited effect on position sense as measured by the arm position matching task

To assess position sense in the arm position matching task, participants were required to move one arm to a location that matches the location of the other arm (the right arm on Fig.8A and B). The latter was passively moved to one of four possibles locations by a robotic manipulandum (the left arm on Fig.8A and B). Participants could readily perform that task as illustrated on Fig.8A for a young participant and on Fig.8B for an older one. Matching errors were present for most targets as the participants did not perfectly match the location of their hands. The distance between the location of the passively moved hand and the matching hand was quantified as absolute position error. In addition, while the errors were quite variable across targets, these errors were quite consistent for the different trials of each target, which we quantified as the inter-trial variability.

**Figure 8.**
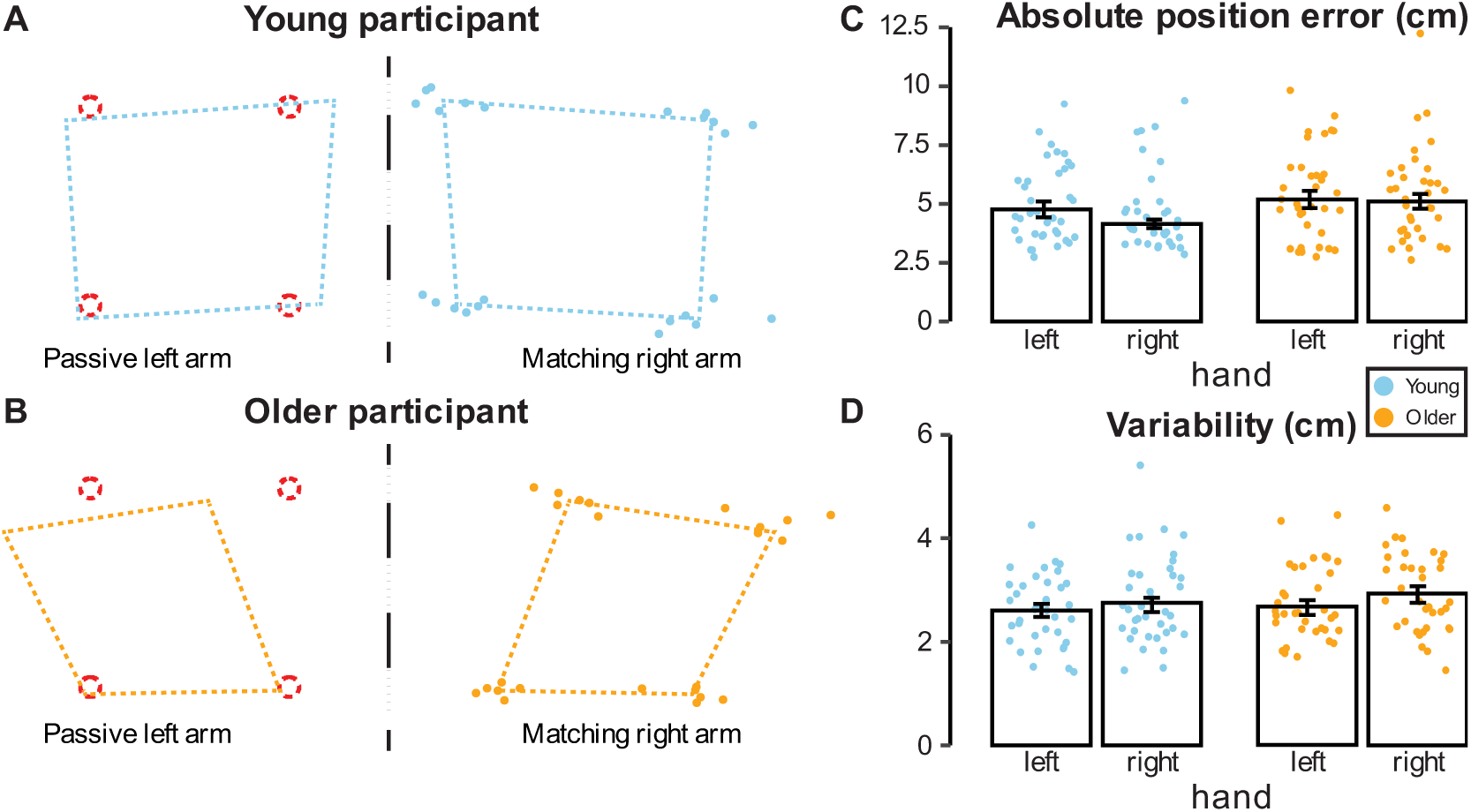
Arm position matching task. Left arm assessment of a typical young (A) and older (B) participant. The left arm of the participant was passively moved by the robot to a target location (dashed red circles), the participant actively mirror-matched with the right arm. Solid dots represent individual matches with the right arm. Dashed colored lines represent mean matches and are mirrored to illustrate how the matches map onto the original targets. The vertical dashed line represents the body-midline which is the reference for mirroring. C) Results for absolute position error and D) inter-trial variability. Dots represent mean parameter values of individual participants. The height of the bars represents group means, the error bars represent the standard error of the mean.

In our sample, the mean absolute position error ranged from 2.62cm to 12.24cm across age groups (Fig.8C). Across both hands, the young adults (M = 4.45 ± 0.23 cm) had smaller errors than the older adults (5.15 ± 0.23cm, main effect of age on absolute position error: *F*(1,36.1) = 4.26, *p* = 0.046, *ES* = 0.39, *CI95(ES)* = [-0.08, 0.86]). We found no evidence for a difference in performance between the left (M = 4.97 ± 0.24cm) and the right hand (M = 4.60 ± 0.22; main effect of hand on absolute position error: *F*(1,44) = 1.49, *p* = 0.23, *ES* = 0.20, *CI95(ES)* = [-0.14, 0.59]). We also found no evidence that performance across hands was influenced by age (interaction effect of hand and age on absolute position error, *F*(1,44) = 0.86, *p* = 0.36).

The borderline significance of the main effect of age could be due to an absence of effect or to a lack of power. To dissociate between these two possibilities, we ran a meta-analysis with data from this study and five other studies conducted in our laboratory (Fig.9). This meta-analysis provides evidence for smaller absolute position error in arm position matching in young adults compared to older adults (*pooled common effect estimate* (Cohen’s d): 0.25, *95CI* = [0.06, 0.44], p=0.01). However, the effect size of this difference was small (d<0.3). One would need a group of 253 young and 253 older participants to detect such a difference with 80% power, which exceeds the typical sample size of proprioception studies. Furthermore, with such effect size, the superiority index is only 57%, meaning that, if one were to test two random participants (one young, one old), the absolute position error of the young participant would only be lower than that of the older participant in 57% of the cases.

**Figure 9.**
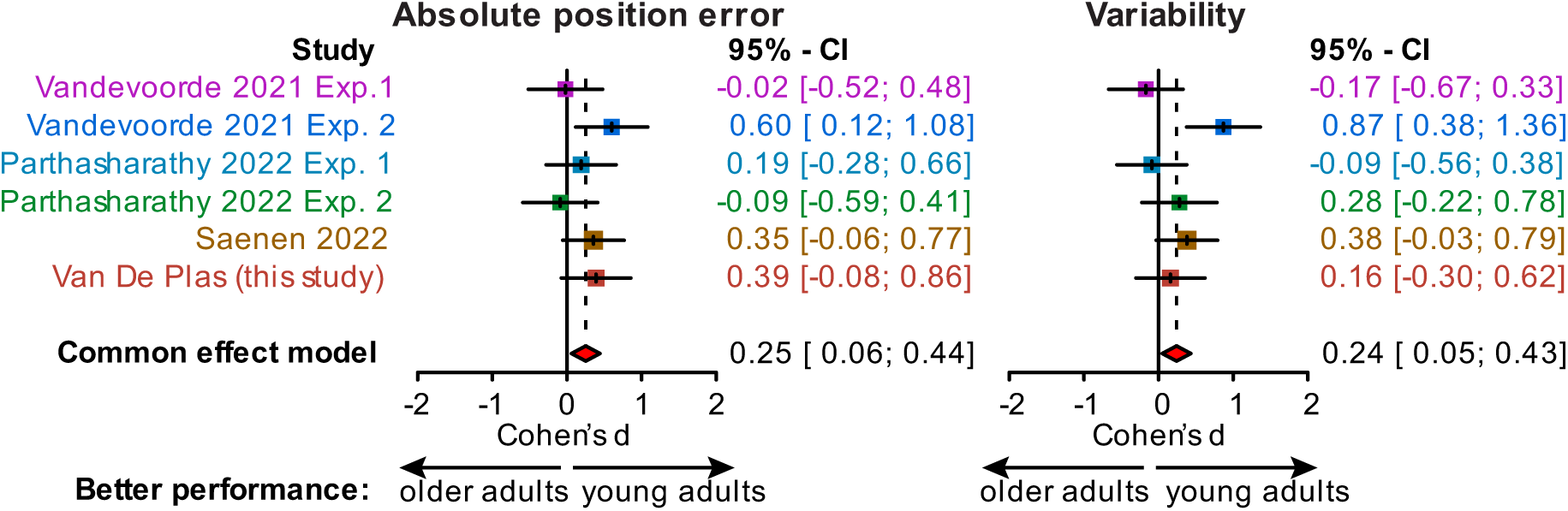
Meta-analysis for the absolute position error (left panel) and for the variability (right panel) in the arm position matching task. The six included studies were conducted in our laboratory. Red diamonds denote average effect size (Cohen’s d) estimated with common effect model.

Next to the accuracy of the matching, we also wanted to assess the consistency, which we inferred from the inter-trial variability of the matched positions (Fig.8D). At the group level, young adults (M = 2.66 ± 0.09cm) and older adults (M = 2.79 ± 0.11cm) were equally consistent in their hand position matches. Across age groups, consistency of hand positions had similar values (left hand: M = 2.63 ± 0.10; right hand: M = 2.81 ± 0.11). We found no evidence that age or the hand used influenced the consistency of the participants (main effect of age on inter-trial variability: *F*(1,43.3) = 0.70, *p* = 0.41, *ES* = 0.16, *CI95(ES)* = [-0.14, 0.49]; main effect of hand on inter-trial variability: *F*(1,42.2) = 1.95, *p* = 0.17, *ES* = −0.21, *CI95(ES)* = [-0.57, 0.10]). There was also no evidence that the consistency for both hands depended on age (interaction effect of age and hand on inter-trial variability: *F*(1,42.2) = 0.35, *p* = 0.56).

Again, to dissociate between an absence of effect and a lack of power, we ran a meta-analysis with data from this study and five other studies conducted in our laboratory (Fig.9). This meta-analysis provides evidence for higher consistency in arm position matching in young adults compared to older adults (*pooled common effect estimate* = 0.24, *95CI* = [0.05, 0.43], p=0.014). As for the absolute position error, the effect size of this difference was small (d<0.3). One would need 275 participants per age group to detect such a difference with 80% power. In this case, the superiority index would be only 56.7%, suggesting that the variability outcome largely overlaps between the two age groups.

#### Older adults are as sensitive to angular deviation from a planned movement as young adults but exhibit more biases

In the perceptual boundary task, participants made reaching movements towards a target during which their hand was deviated away. The goal of the participants was to determine in which direction their hand was deviated. The size of the deviation was adapted in direction and in size in function of the ability of the participants to correctly identify the direction of the perturbation (adaptive PEST procedure, Fig.10A). In the example from Fig.10A, the participant could accurately indicate on which side of the target s/he ended up with the exception of trial 4 where the angular deviation was 0.5° to the left but the participant judged it to be to the right. After completing eight blocks like this, a psychometric curve was fitted to the data. The obtained psychometric curve (Fig.10B) shows that this participant was able to judge to which side of the target s/he reached to when the angular deviation was large (±25° or ±23.5°) as this were respectively associated with 100% of correct responses (represented by the blue dots in Fig.10B). The participant was never answering “to the right” when the angular deviation was more than 20deg towards the left and answering “to the right” in 100% of the trials when the angular deviation was more than 20deg towards the right. In contrast, for angular displacement around −5°, the participant was more uncertain and answered almost randomly (probability of saying right or left for a given displacement is close to 50%). The gradual change in the perception of the direction of the proprioceptive deviation can be represented by the psychometric curve. Interestingly, this participant had a leftward bias and subjectively considered that s/he was reaching straight to the target (where the psychometric curve is equal to 0.5) when the hand was deviated 5° to the left of the target (inflection point). In addition, the sensitivity of the participants to the angular deviations of the hand trajectory can be represented by the slope of the psychometric curve between the 0.25 and 0.75 range, which corresponds to the uncertainty range. The higher the slope is, the smaller the uncertainty range is.

**Figure 10.**
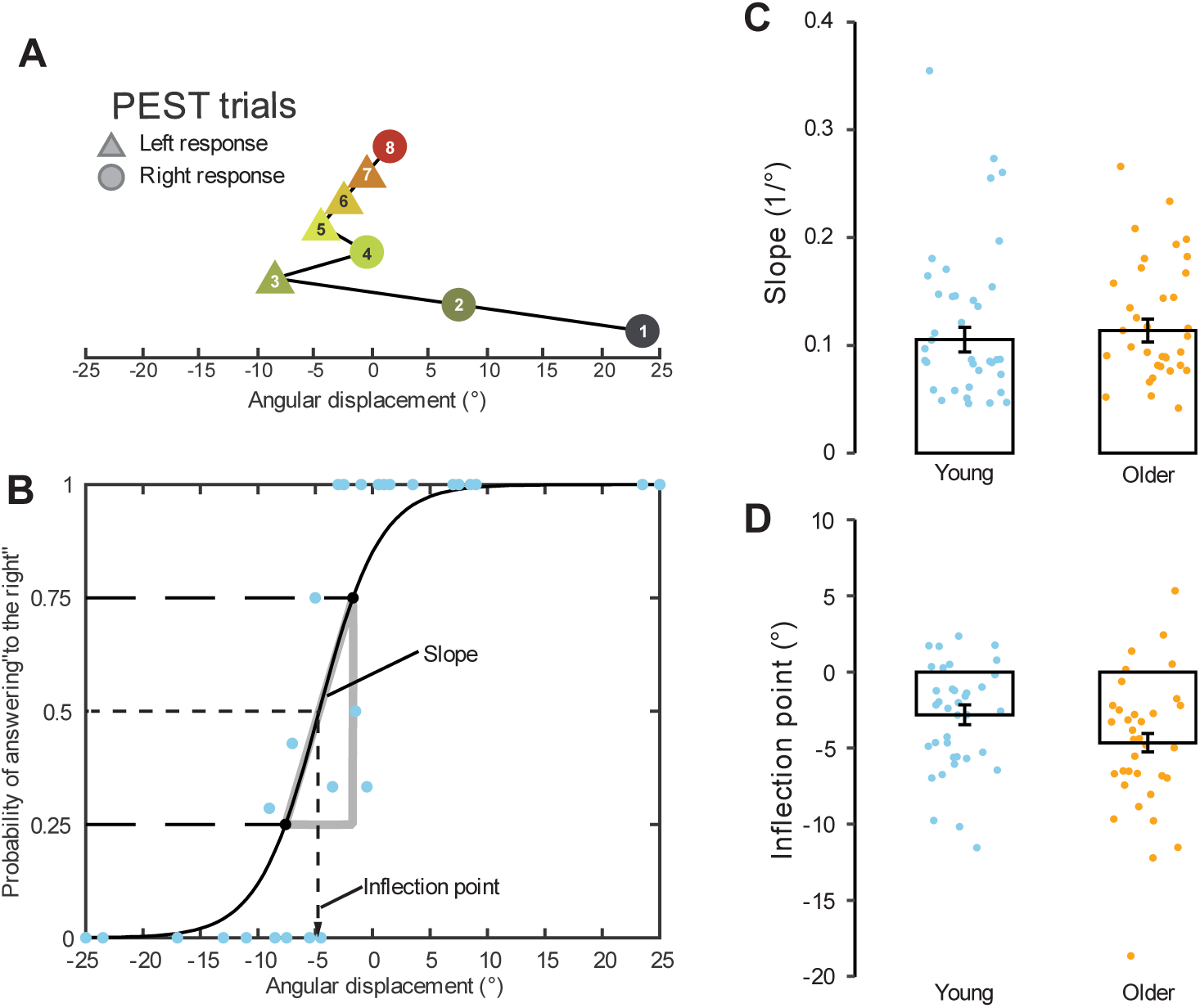
Perceptual boundary task. A) Angular deviation for each trial in block one of a young example participant. The number of trials in a block depends on the performance according to the adaptive parameter estimation by sequential testing (PEST) procedure. B) Answers from one young participant represented as the probability of answering right as a function of angular displacement. A psychometric curve is fitted from which the slope and inflection point can be computed. C) Slope results. Dots represent mean parameter values of individual participants. The height of the bar represents group mean; the error bars represent the standard error of the mean. D) Results for inflection point. Dots represent mean parameter values of individual participants. The height of the bars represents group means, the error bars represent the standard error of the mean.

To quantify proprioceptive accuracy as measured by the perceptual boundary task in our sample, the slope and the inflection point of the psychometric curve were calculated for each participant (Fig.10C and D). The slope is an estimate of how sensitive to the proprioceptive angular deviations participants were. Young (M = 0.105 ± 0.011) and older (M = 0.114 ± 0.011) adults were equally sensitive to angular displacements from the target. We found no evidence for an age-related difference (Fig.10C, effect of age on slope of the psychometric curve: *t*(42) = 0.55, *p* = 0.59, *ES* = −0.14, *CI95(ES)* = [-0.68, 0.32]). To further interpret this absence of significance, we conducted a meta-analysis based on this study and four other studies using this task. The meta-analysis confirms our findings that there is no evidence for an effect of age on slope over these five studies based on data from 162 young and 167 older adults (Fig.11, *pooled common effect estimate* = −0.09, *95CI* = [-0.31, 0.13]). If anything, older adults were more sensitive to the lateral displacement (higher slope) than younger ones. In this case, the superiority index is only 47.4%, which means that, if one were to test one young and one older participant, the slope of the young participants would be steeper than that of the older adults in only 47% of the cases.

**Figure 11.**
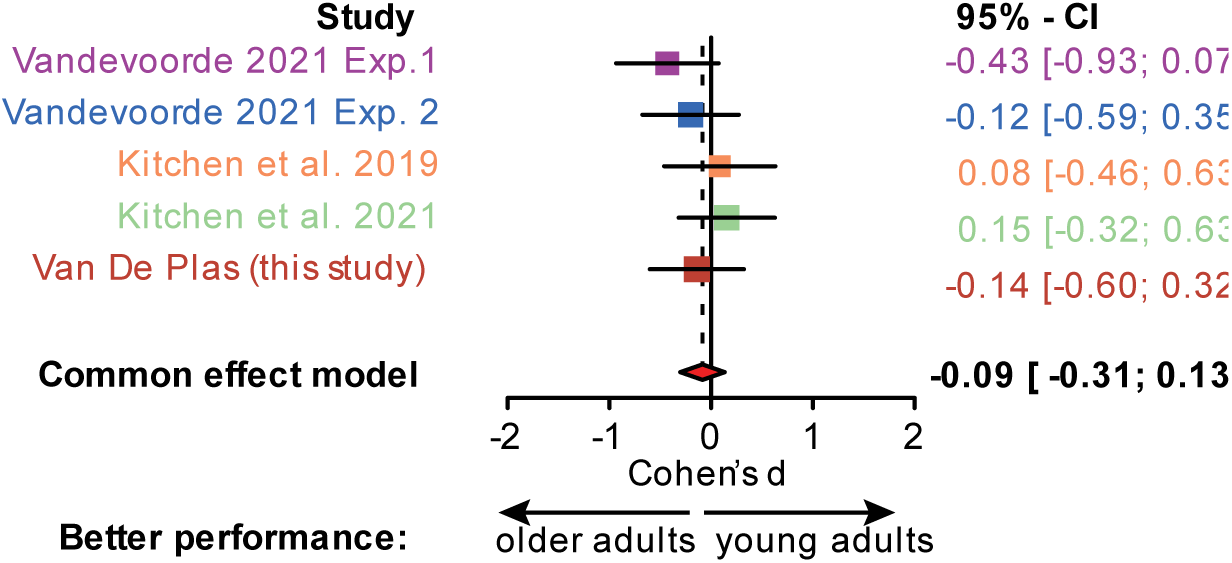
Meta-analysis for slope of the perceptual boundary task. Red diamond denotes average effect size (Cohen’s d) estimated with common effect model.

Next to discriminative sensitivity to angular displacements, the inflection point indicated what displacement felt like zero displacement from target during reaching, i.e. reaching straight ahead. At the group level, an angular displacement to the left felt like reaching straight ahead for young (M = - 2.82 ± 0.65°) and for older adults (M = −4.65 ± 0.61°). We found evidence for an effect of age (Fig.10D, effect of age on inflection point of the psychometric curve: *t*(41.96) = 2.08, *p* = 0.04, *ES* = 0.52, *CI95(ES)* = [0.02, 1.02]) indicating that older adults were more biased in their subjective feeling of reaching straight ahead compared to young adults.

#### Haptic perception is better in young compared to older adults

So far, we assessed proprioceptive accuracy relative to single-point targets in the arm position matching task and the perceptual boundary task. We also wanted to look at more complex targets representing more than one point in space. In the shape reproduction task, targets are geometrical shapes that are first explored by passive movement, and later reproduced by active movement. A typical young participant reproduces a pentagon quite accurately while the shape produced by a typical older adult matches the target shape less accurately (Fig.12A).

One way to quantify the similarity of the shapes is the cross-correlation between the X or Y position of the left hand during reproduction and that of the right hand during haptic exploration (Fig.12C). On average, the cross-correlation between the explored and the reproduced shapes was higher in young (M = 0.89 ± 0.005) compared to older adults (M = 0.86 ± 0.005) for both X and Y direction (main effect of age: *F*(1, 42.92) = 12.31, *p* = 0.001, *ES* = 0.77, *CI95(ES)* = [0.43, 1.13]). Across age groups, the cross-correlation between the explored and the reproduced shapes was higher in the Y direction (M = 0.88 ± 0.005) than in the X direction (M = 0.87 ± 0.004, main effect of direction: *F*(1, 43.82) = 6.62, *p* = 0.01, *ES* = −0.27, *CI95(ES)* = [-0.64, 0.06]). We found no evidence that the effect of age on the cross-correlation differed between the X and Y directions (*F*(1, 43.82) = 1.77, *p* = 0.19).

Furthermore, the distance between the explored and reproduced shapes after optimal superimposition was, on average, smaller for young adults (M = 0.22 ± 0.01) compared to older adults (M = 0.25 ± 0.008) (Fig. 12C, Procrustes analysis, *t*(39.52) = 2.48, *p* = 0.017, *ES* = −0.61, *CI95(ES)* = [-1.14, −0.12]) providing again evidence for worse shape reproduction in the older group of participants. Next to reproduction, shape discrimination was also impaired in older adults. On average, young adults (Fig.12D, M = 77.10 ± 2.62) could correctly identify more shapes than older adults (M = 68.75 ± 3.69) but this effect did not reach significance (*t*(35.96) = 1.85, *p* = 0.073, *ES* = 0.47, *CI95(ES)* = [0.03, 0.93]).

**Figure 12.**
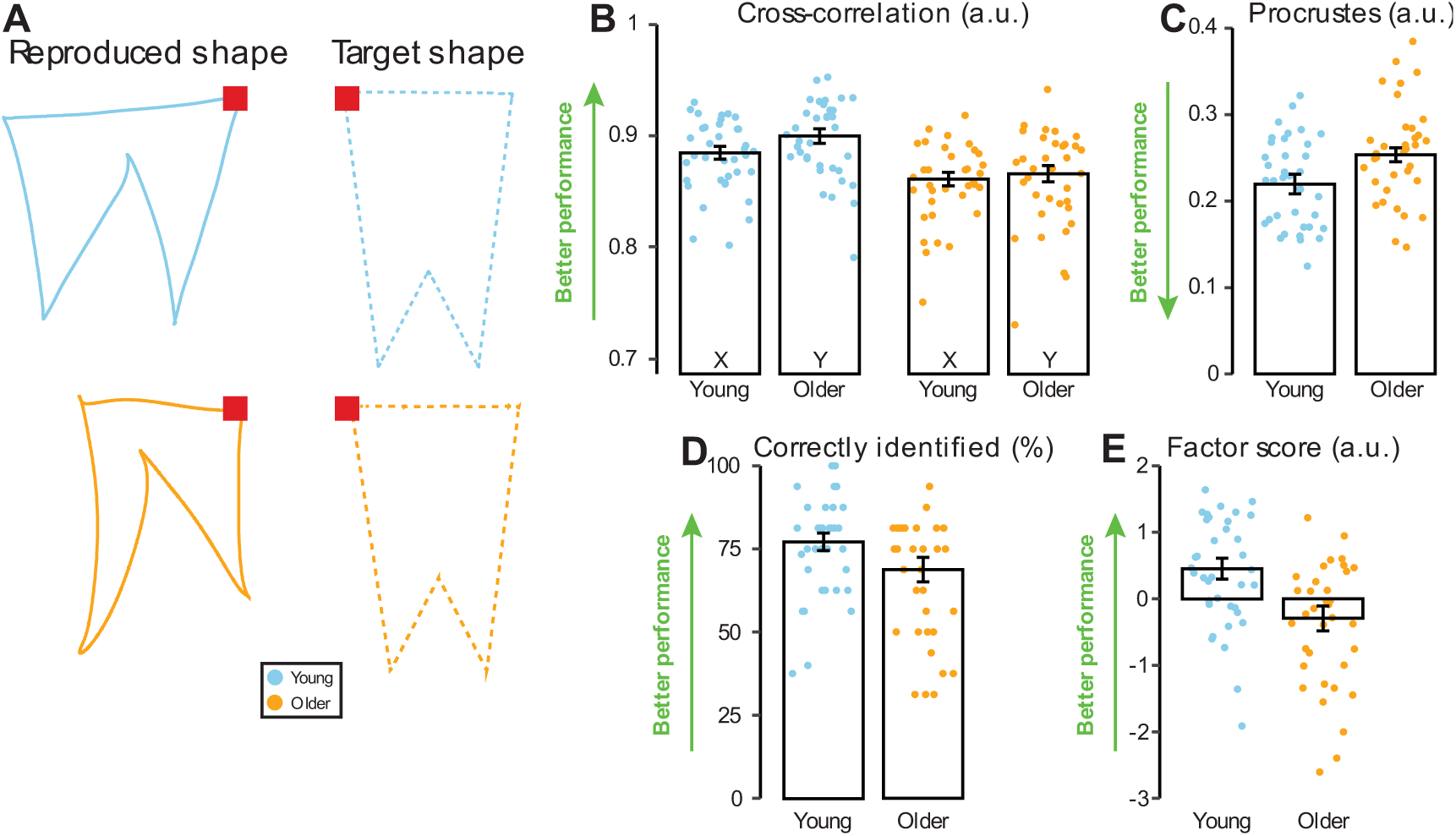
Shape reproduction task. A) Example data of one trial for a typical young and older participant. The shape is first passively explored with the right hand (step 1). Then, the participant actively reproduces the shape with the left hand (step 2). The right and left red square are the starting and end points for exploration and reproduction, respectively. B) Cross-correlation in the X- and Y direction. C) Procrustes analysis. D) Percentage correctly identified. E) Factor score. Better or worse performance is indicated for each parameter. Dots represent mean parameter values of individual participants. The height of the bars represent group means, the error bars represent the standard error of the mean.

To provide a global view of the effect of aging on this haptic perception task, we performed a factor analysis based on the four parameters discussed above. On average, young adults (Fig.12E, M = 0.45 ± 0.16) had a better performance (as quantified by the factor score) compared to older adults (M = −0.29 ± 0.19, *t*(39.41) = 3.08, *p* = 0.004, *ES* = 0.78, *CI95(ES)* = [0.30, 1.35]). In this case, the superiority index amounts to 0.72, meaning that the factor score of a young adult would be higher than that of an older adult in almost three quarters of the cases.

Given that this task requires a lot of working memory because the participants had to remember the explored shape, we tested the influence of working memory capacity on the performance of the participants. Indeed, we found a medium correlation between the factor score and visual-spatial working memory capacity (multilevel correlation, *r* = 0.29, *p* = 0.016). Yet, even after controlling for visual-spatial working memory capacity, there’s evidence for an age-related effect on the shape reproduction factor score (*F*(2,68) = 9.704, *p* < 0.001, d = 1.06).

### Integration/recalibration of proprioception with vision

Proprioception contributes to the estimation of hand position together with other senses. For instance, in the presence of vision, both modalities (vision and proprioception) are used to obtain an estimate of hand position via multisensory integration. Multisensory integration follows Bayesian inference implying that the contribution of each modality is weighted based on the reliability of its signal. Therefore, by manipulating visual feedback, we can make inferences about the relative weight of proprioceptive input for state estimation, hence measuring the relative weighting of vision and proprioception and thus, indirectly, the reliability of proprioceptive information. In the next sections, we look at the influence of perturbed visual feedback on estimates of hand position in a visuomotor adaptation task and a multisensory integration task.

#### No evidence for an age-related difference in level of implicit adaptation in the task-irrelevant clamped feedback task

In the task-irrelevant clamped feedback task, sensorimotor recalibration is induced by uncoupling the direction of the cursor motion from that of the hand, hence creating a mismatch between proprioceptive and visual information. According to Tsay et al. (2022), the extent of the recalibration due to this mismatch is directly related to the reliability of the proprioceptive signal. In all participants, the imposed mismatch between the motion of the cursor and of the hand induced a drift away from the direction of the perturbation as shown for a typical young and a typical older participant (Fig.13A). By the end of the adaptation phase, both participants reached around 25° to the right of the target in response to the leftward perturbation. At the population level, we found no evidence for an age-related difference in the amount of implicit adaptation (Fig.13B). To quantify adaptation, we computed the mean hand angle of the last 30 trials in the test phase (shaded area in Fig.13A and B) corrected for baseline (Fig.13C, M of young adults = 20.54 ± 1.14; M of older adults = 20.83 ± 1.44; *t*(38.31) = 0.16, *p* = 0.87, *ES* = −0.04, *CI95(ES)* = [-0.58, 0.44]).

**Figure 13.**
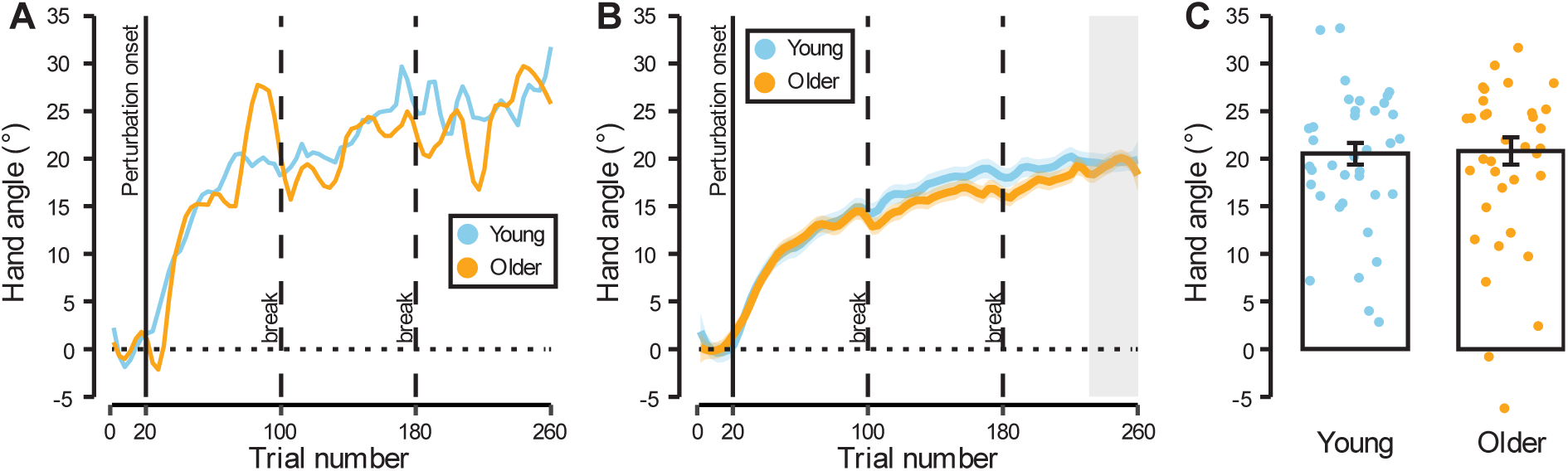
Task-irrelevant clamped feedback task. A) Example data of a typical young and older participant. B) Data of all participants averaged by group. Solid curves represent the group mean of the hand angle for each trial. The areas around the curves represent the standard error of the mean. Clamp onset is indicated by a solid vertical line. Dashed vertical lines indicate 1 minute breaks. C) Mean hand angle of the last 30 trials (grey shaded area in A and B). Dots represent individual mean hand angles. The height of the bars represents the group mean and the error bars represent the standard error of the mean.

As always, the absence of significance can be due to a lack of power or an absence of effect. To distinguish between these two possibilities, we performed a meta-analysis based on six studies (including this one) that all used the task-irrelevant clamped feedback task in young and older adults. The results of the meta-analysis show that older adults exhibit more implicit adaptation than young adults (Fig.14, *pooled common effect estimate* = −0.34, *95CI* = [-0.52, −0.16]), but the effect size is small and requires around 140 participants per group to show the effect with 80% power. In this case, it is fare more likely to have higher adaptation in older participants than in younger ones as the superiority index only amounts to 40% for young adults.

**Figure 14.**
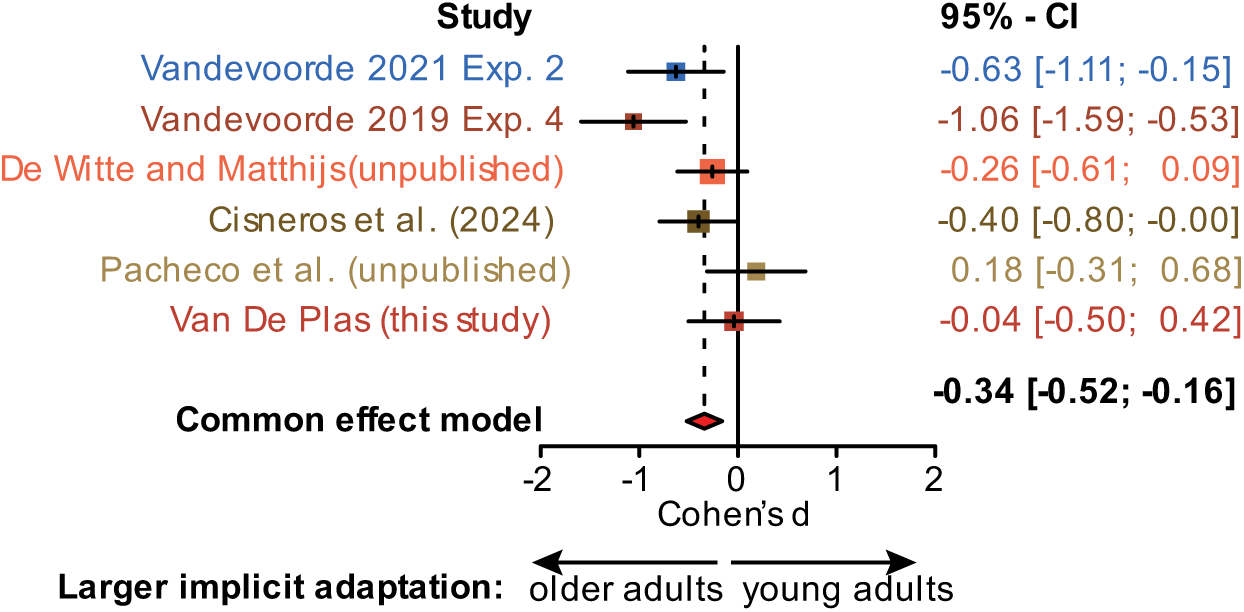
Meta-analysis for implicit adaptation of the task-irrelevant clamped feedback task. Red diamond denotes average effect size (Cohen’s d) estimated with common effect model.

#### Age-related difference in the weighting of proprioceptive and visual information in an integration task

Besides looking at the effect of continuous perturbation on reaching movement over many trials, we wanted to look at the effect of perturbation in single trials. The influence of perturbation on movement within a single trial can give an indication of reliability of vision and proprioception to estimate the position of the hand. High reliance on proprioception would lead to a smaller effect of a visual perturbation on the estimate of hand position.

In the integration task, the participant makes a movement to a remembered target with visual feedback (dotted traces in Fig.15A). Then, the cursor disappears, and the participant must perform the return movement to the starting position in the absence of any visual information about the location of his/her hand and of the starting position (solid black traces in Fig.15A). In some trials, the direction of the cursor is deviated away from the actual hand motion by either 15° or 25° (colored traces in Fig.15A). This causes a mismatch between visual and proprioceptive information about the hand location, hence influences the estimate of the hand position. To return to the target, participants use this estimate in order to execute an accurate return movement. Therefore, the direction of the return movement (solid traces in Fig.15A) provides information about the weight of proprioceptive information on the estimated hand location. For both young and older adults, the perturbed visual feedback influences the direction of the return movement (Fig.15A).

For each trial, from each participant, we obtained the direction of the return movement (blue and orange dots in Fig.15B and C). In the absence of any perturbation (75% of the trials in this experiment, black traces on Fig. 15A), the return movement were quite accurate (0° in Fig. 15B and C) but variable. We considered this variability was linked to the consistency of the proprioceptive estimates (Fig. 15F). Consistency was calculated as the standard deviation of initial direction error of return movement for all trials without perturbation (0° rotation). On average, we found no evidence for an age-related difference in the consistency (Fig.15F, M of young adults = 5.02 ± 0.33, M of older adults = 7.71 ± 1.55, t(21.69) = 1.69, p = 0.11, ES = 0.44, CI95(ES) = [0.06,0.89]).

**Figure 15.**
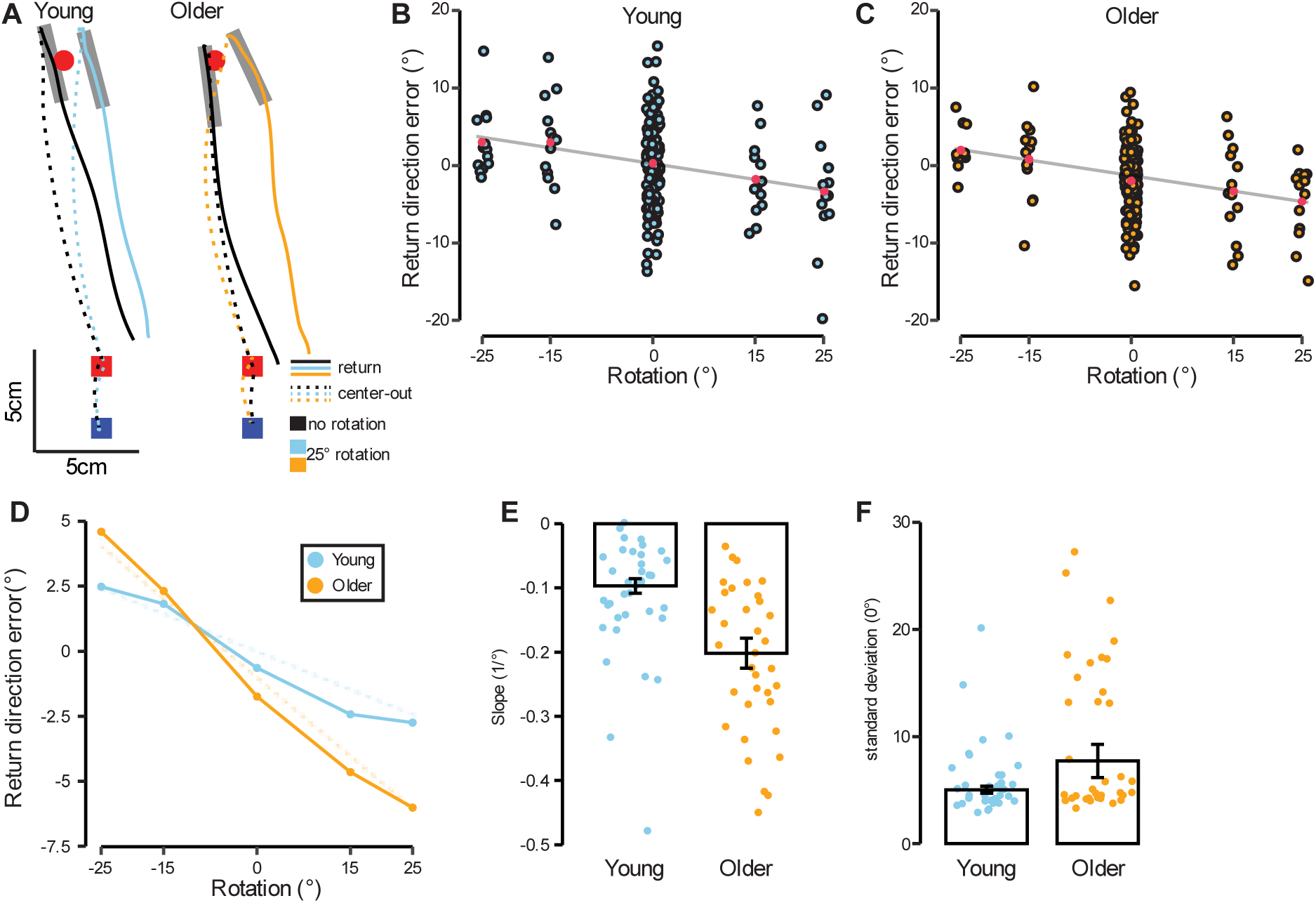
Integration of proprioception and vision task. A) Example data of a young (left) and older (right) participant with the same memorized target (red circle). The blue square represents the pre-movement target and the red square the starting target. Dashed lines represent the center-out reaching movement and the solid lines the return movement. Grey bars represent the initial direction of the return movement. The black lines are for the no-rotation condition and the colored ones for a 25° rotation to the right. B and C) Individual trial data (each dot) for the two participants (young participant B, older participant C) from panel A. The red dots represent the mean per rotation angle. The grey line represents the linear fit on the means. D) Average direction error for the return movement per rotation angle for both groups. The dashed lines represent the linear fit. E) Comparison of the slope of the linear fits (as illustrated in panels B and C) for both age groups. Each dot represents the slope of one participant, the height of the white bar represents the group mean and the error bar the standard errors of the group mean. F) Comparison of the accuracy of return movement in the 0° rotation condition. Each dot represents the standard deviation of one participant, the height of the white bar represents the group mean and the error bar the standard errors of the group mean.

In the presence of perturbation (±15 and ±25°), the return movements were consistently biased away from the starting target. When deviated towards the left (negative rotations in Fig.15B-D), the return movements were biased towards the right (positive initial direction error in Fig.15B-D) and vice-versa. At the group level, the impact of visual perturbation was larger for older adults than for young ones (Fig. 15E).

To quantify the sensitivity of the direction of the return movement to the visual perturbation, we fitted a regression line from these five points. For each participant, the slope of the regression line indicates how much the direction of the return movement was influenced by the visual perturbation. The slope is thus an indirect measure of the reliability of proprioceptive information. The steeper the slope, the lower the reliability of the proprioceptive information and the higher the weight of visual information. On average, the slope of older adults (M = −0.20 ± 0.02) was steeper compared to that of young adults (Fig.15E, M = −0.10 ± 0.01). We found evidence for a significant age-related difference with a large effect size (Fig. 15E, t(28.5) = 4.02, *p* < 0.001, *ES* = 1.04, *CI95(ES)* = [0.55, 1.71]) and a superiority index of 77%. This analysis confirmed that the rotated visual information in this task had a larger effect on the estimated hand position of older adults compared to young adults, which suggests that the relative weight of proprioception to the estimation of hand location is lower in older adults. Following Bayes rules, this would suggest that the reliability of proprioceptive information is lower in older adults.

#### Tactile detection threshold is lower (better) in young adults compared to older adults

Given the influence of touch on position sense (Delhaye et al., 2018; Kuling et al., 2016; Proske & Gandevia, 2012; Zopf et al., 2011), we also quantified touch acuity at the index fingertip with the Bumps test. In this task, participants had to indicate which plate out of five plates had a small bump in the middle. The height of the bumps varied between 5 µm to 25 µm by steps of 5 µm. On average, young adults detected smaller bumps than older adults. For instance, only two older participants (5.71%) reliably detected the 5µm bump while 19 younger participants (51.35%) did. As a result, on average, young participants had a significantly lower threshold for the detection of bumps (M = 7.39 ± 0.69 µm) than older participants (Fig.16, M = 11.43 ± 1.20 µm, *t*(31.64) = 2.91, *p* = 0.006, ES = 0.75, *CI95(ES)* = [0.53, 1.30], superiority index: 70%).

**Figure 16.**
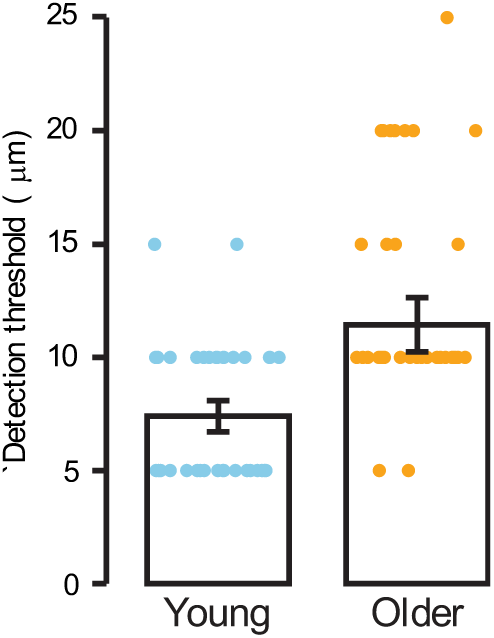
Bumps task. Each dot represents the detection threshold of an individual participant. The height of the bars represents the group mean and, the error bar the standard error of the mean.

### The effect of age on motor function is task-dependent

For the first motor task, participants had to reach to different targets. For this simple motor task, young and older participants exhibited similar performance (Fig.17A and B). Movement trajectories of both participants were smooth, without much overshooting of the target.

In the visually guided reaching task, visuomotor capabilities were quantified by two factor scores. One represented motor control, the other represented motor speed (Saenen et al., 2023). On average, young and older adults had similar factor scores for the score related to motor control (Fig.17C, YA: 0.06 ± 0.18; OA: −0.17 ± 0.17; *t*(41.96) = 0.93, *p* = 0.36, *ES* = 0.23, *CI95(ES)* = [-0.25, 0.78], superiority index: 56.4%) and for the one related to speed (Fig. 17D; YA: = −0.24 ± 0.16; OA: 0.11 ± 0.16; *t*(41.94) = 1.56, *p* = 0.13, *ES* = 0.39, *CI95(ES)* = [-0.10, 1.03]).

**Figure 17.**
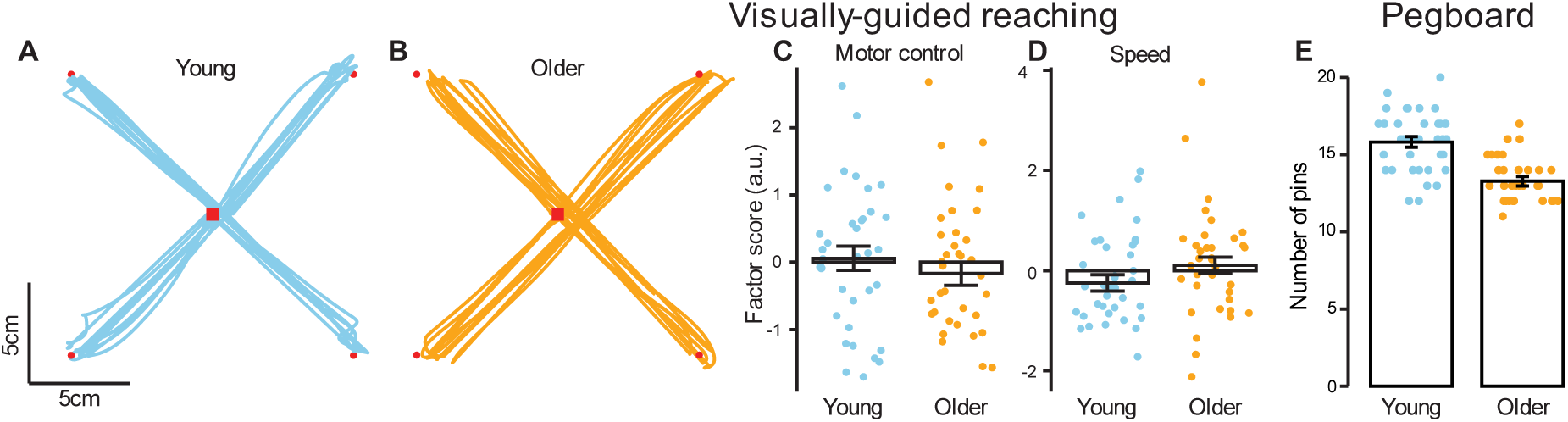
Motor tasks: Visually guided reaching task (A-D) and Purdue pegboard task (E). A and B) Center-out movements for young participant (A) or older participant (B), reaching from the central target (red square) to the peripheral targets (red dots). C and D) Comparison of the factor score for “motor control” (C) and for “speed” (D) across age groups E) Comparison of the number of pins placed in 30s during the Purdue pegboard task. In panel C-E, each dot represents the mean value of one participant. Bar height represents the group mean and the error bar the standard error of the mean.

In contrast, in the Purdue pegboard task where participants placed pins in the holes of a wooden board as fast as possible, young adults (M = 15.83 ± 0.34) outperformed older adults (Fig. 17E, M = 13.29 ± 0.32). This motor task exhibits the largest effect size of our task battery in terms of age-related difference (*t*(41.97) = 5.47, *p* < 0.001, *ES* = 1.37, *CI95(ES)* = [0.91, 2.32]).

### Clear age-related difference in spatial working memory capacity

Finally, we tested the ability of participants to remember the location of dots among a set of 16 rectangular targets and inferred working memory capacity from their behavior. Data of one young participant was excluded from this analysis because of inconsistencies in the answer pattern resulting in a K=0. Young adults (M = 4.43 ± 0.16) exhibited larger working memory capacity than older adults (Fig.18A, M = 3.62 ± 0.14, *t*(40.79) = 3.88, *p* < 0.001, *ES* = 0.98, *CI95(ES)* = [0.32, 1.53]). This higher working memory capacity was also reflected in a higher percentage of correct answers in young (M = 92.99 ± 0.90) than older adults (Fig. 18B, M = 86.31 ± 1.41; *t*(33.77) = 3.97, *p* < 0.001, *ES* = 1.02, *CI95(ES)* = [0.51, 1.61]).

**Figure 18.**
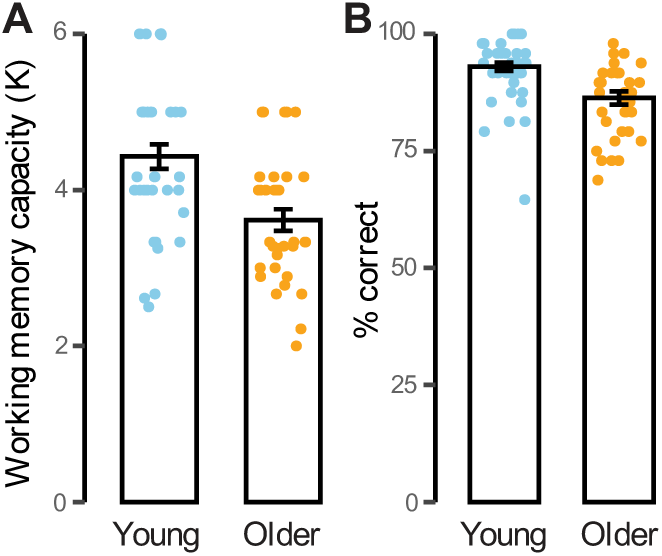
Spatial working memory task. A) Comparison of working memory capacity K, and B) the percentage of correct answers. Each dot represents the mean value of one participant. Bar height represents the group mean and the error bar the standard error of the mean.

To check whether performance on proprioceptive tasks was influenced by differences in visual-spatial working memory, we calculated multilevel correlations (Fig.19). The haptic proprioception task is the only task showing a significant correlation with visual-spatial working memory, hence we controlled for this confounding factor in the analysis of this task (cfr supra).

**Figure 19.**
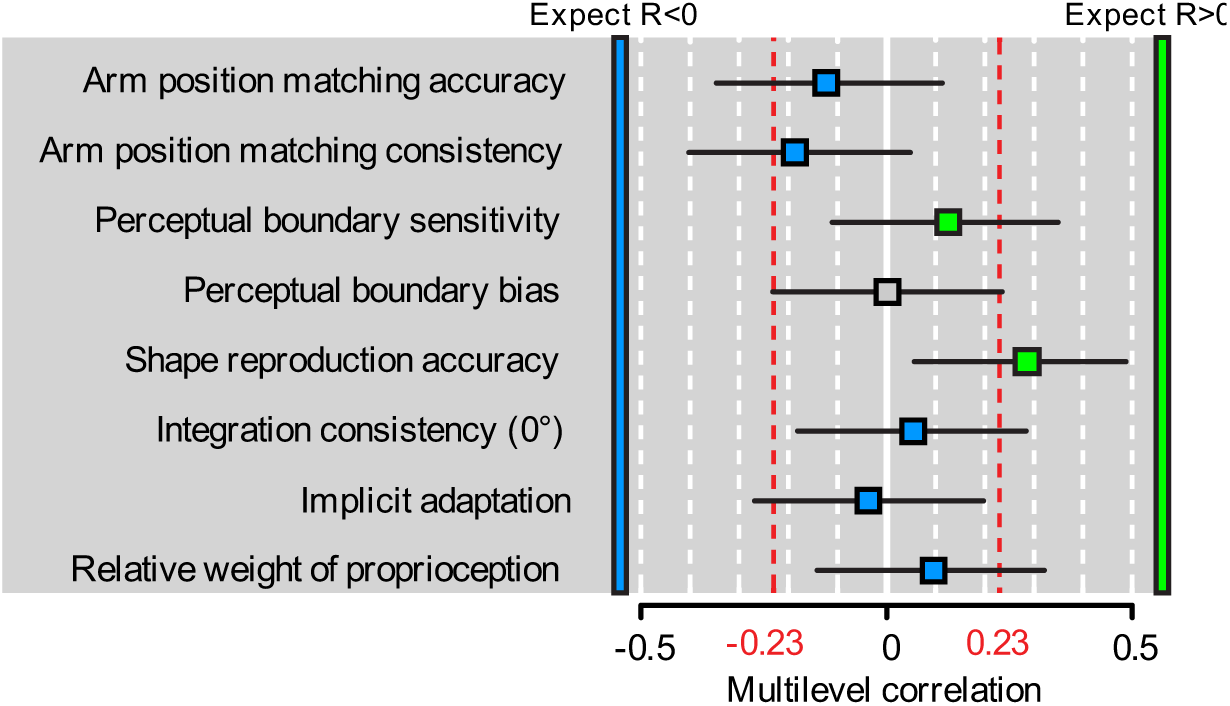
Multilevel correlations of parameters with visuo-spatial working memory capacity. Box color indicates the direction in which correlation was expected (green for positive, blue for negative, grey: no expectation). Red dotted lines provide the critical values for correlations with N=71 (two-tailed). Lower values are all non-significant.

### Age-related effects are outcome-dependent, task-dependent and domain-dependent

The effect sizes for all analyses presented above (Fig.20) suggest that age-related differences are very variable. While the age-related effect in the cognition domain was clear and large (for one task only), effect sizes in all other domains were very variable, ranging from very small to large. For instance, in the motor domain, the Purdue pegboard task exhibited large age-related differences while visually-guided reaching showed small age effects. Similarly, while the integration of vision and proprioception in the two recalibration tasks is supposed to be based on the same mechanism, the sensory integration task exhibited large age-related difference while there was none in the implicit adaptation task. Finally, even within the outcomes of a single task, we found large effect size differences. For instance, within the perceptual boundary task, the bias was worse in older adults, but their sensitivity was not (if anything, it was slightly better in older adults).

**Figure 20.**
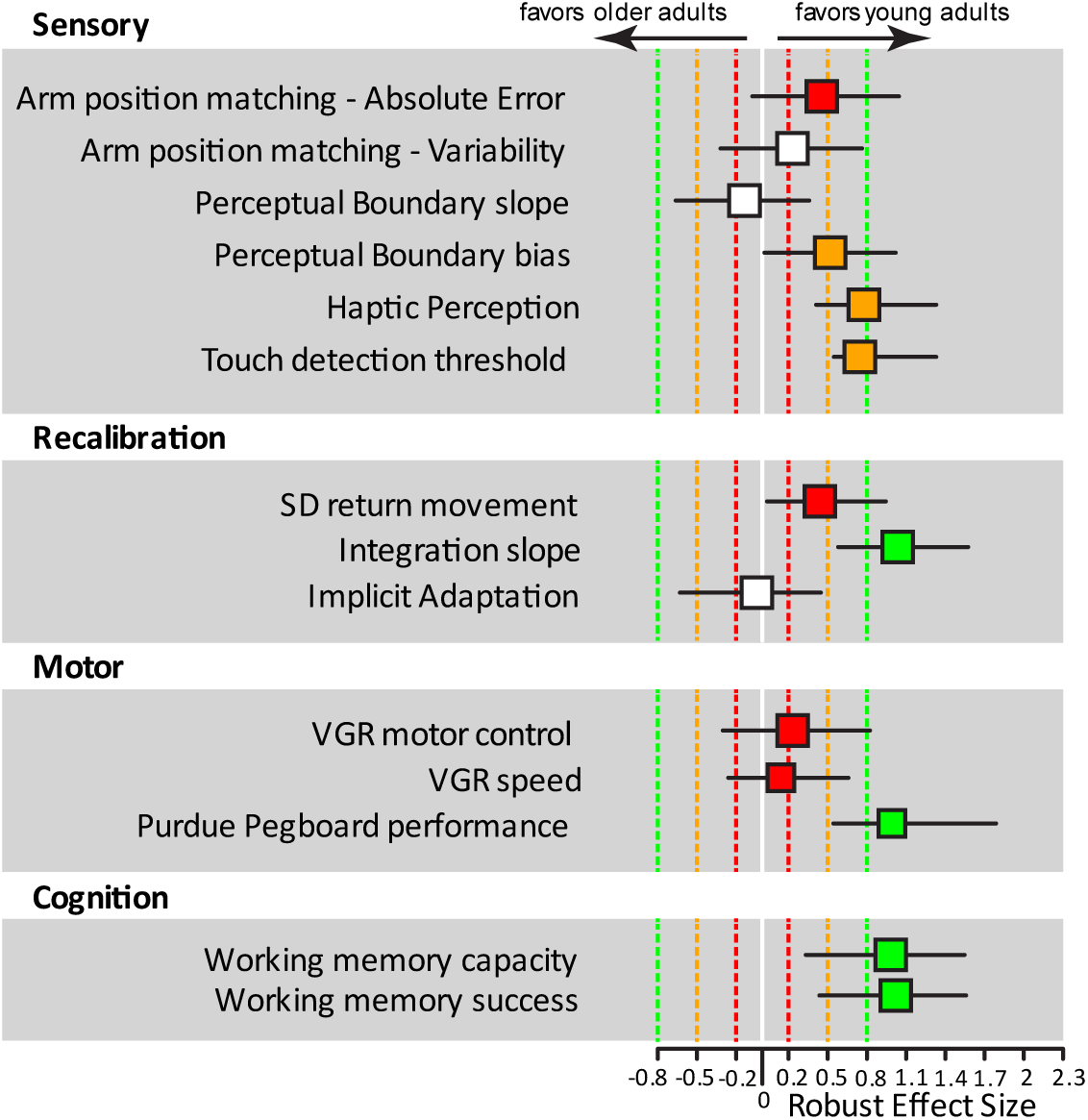
Robust effect size for all parameters obtained from the new sample of participants. The color of the marker corresponds to negligible (white, between 0 and 0.2), small (red, 0.2 to 0.5), medium (orange, 0.5 to 0.8) and large (green, larger than 0.8) in both positive and negative directions. Positive effect sizes indicate better performance by young adults and negative ones by older adults.

This overview suggests that, while we believe that all the proprioceptive tasks and their outcomes are measures of proprioceptive function, there is very little similarity in the age-related effects. If a latent factor linked to proprioceptive function influenced all these outcomes and if aging influenced this latent factor, we would expect a much more widespread effect of age on these outcomes and less between-task variability.

Furthermore, if a latent factor linked to proprioceptive function was responsible for all these task outcomes, we would expect these different outcomes to correlate with each other. To investigate these correlations, we calculated multilevel correlations (to account for between age group differences, Fig. 21). Given our sample size of N=72, we could detect correlation as small as 0.23 (critical value for df=70, two-tailed). None of the correlations between variables representing proprioceptive accuracy/bias reached that value as they were all small to negligible. There was a small negative relationship between the consistency from the arm position matching task and from the perceptual boundary task (*r* = −0.24, *p* = 0.05), which replicates the negative correlation found between the same two variables in Vandevoorde & Orban de Xivry (2021). Within tasks, correlations between variables were also small to negligible. Overall, most correlations were in the expected direction, but none of them were large.

**Figure 21.**
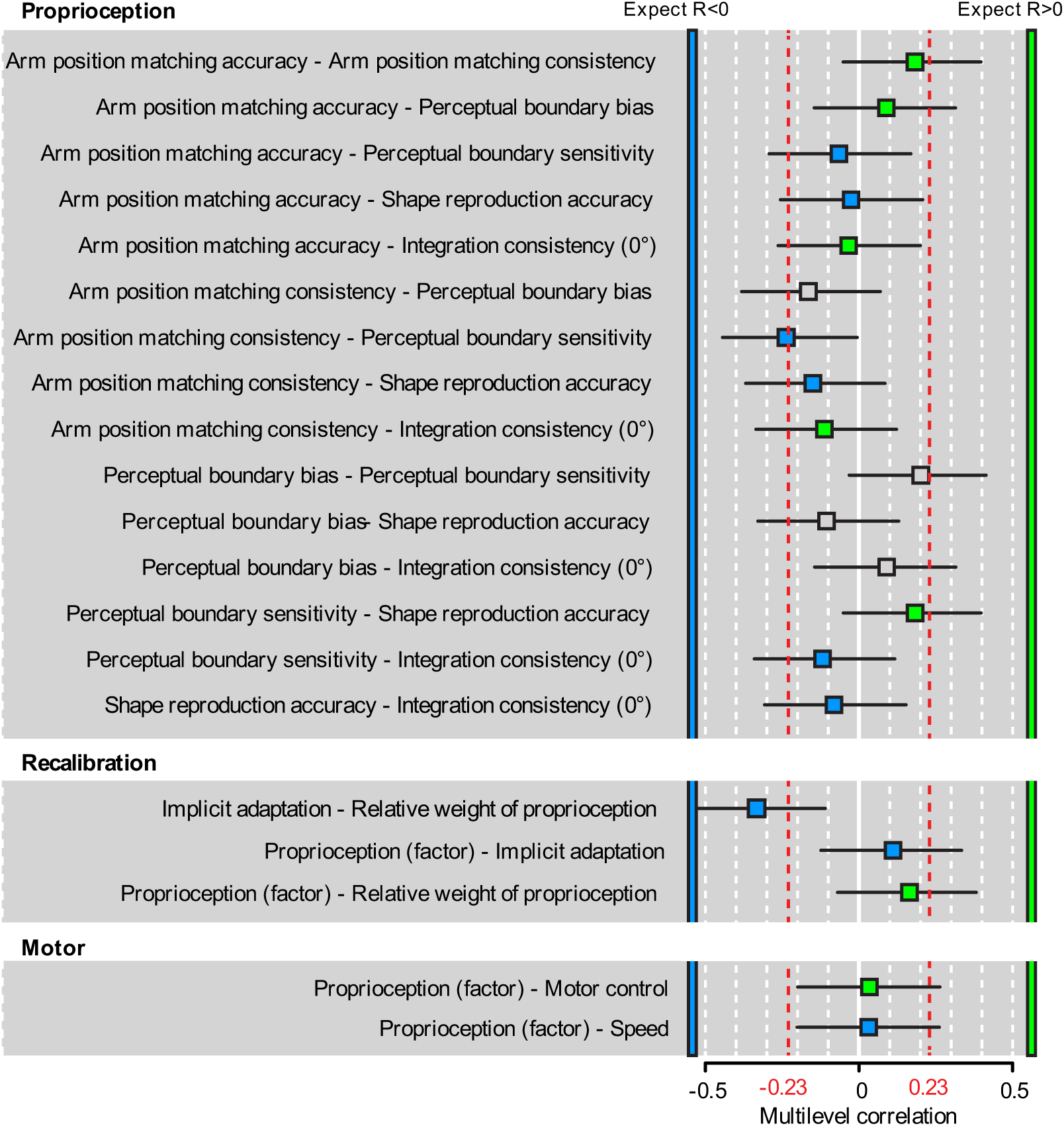
Multilevel correlations between parameters. Box color indicates the direction in which correlation was expected (green for positive correlation, blue for negative correlation, grey: no expectation). Red dotted lines provide the critical values for correlations with N=72 (two-tailed). Lower values are all non-significant.

In the recalibration tasks, we expected that a low reliability of proprioception would lead to higher implicit adaptation and a higher weight of vision in the sensory integration task (i.e. more negative slope). For these reasons, we expected a negative correlation between these two recalibration parameters and indeed found evidence for a medium negative correlation between the two parameters (*r* = −0.33, *p* = 0.004). This correlation was both expected given the similar putative role of the reliability of proprioception in both tasks and surprising because the age effect is very different in both tasks (Fig.20). In addition, despite the putative role of the reliability of proprioception for this task, the correlation between our latent factor linked to proprioception (obtained from a factor analysis on the outcomes of the proprioceptive tasks) and the amount of recalibration found in these tasks was small. Lastly, we did not find any evidence that quality of proprioception is related to motor control or speed.

## Discussion

Textbook knowledge assumes that large age-related declines in proprioception can be found with small sample sizes in any proprioceptive task. Our study demonstrates that the effect of aging on proprioceptive function is not as simple as textbooks often describe. We found that age-related differences in proprioception and in motor function were task-dependent. Despite our relatively large population (N=72 in total), we could not detect significant correlations between the different outcomes linked to proprioception. While age-related effects on proprioception were limited, they were clear for touch, which is another aspect of somatosensory function, and for cognition.

### Age-related differences in proprioception are task-dependent

Depending on the task used, an age-related decline in proprioception is absent even with large sample sizes (e.g. slope of the perceptual boundary task), or a very large sample size is required to reliably find a small effect of age (e.g. variability of the arm position matching task), or a small sample is sufficient to find an effect of age (e.g. influence of vision in the integration task). If aging had a large influence on proprioceptors or on proprioceptive processing, one would expect a large age-related difference in all of these tasks. Yet, we only found a large effect size on a task where proprioceptive reliability was measured indirectly via sensory integration, which opens the possibility that processes other than proprioceptive processing could be impacted by aging such as sensory integration, etc.

Task-dependency is evident in both the sensory (perceptual boundary vs. position matching) and in the recalibration domain (adaptation vs. sensory integration). Within the sensory tasks, arm position matching and perceptual boundary provide a different conclusion on age-related differences. In the meta-analysis of the arm position matching task, both outcome variables provide evidence for a small age-related effect. In the meta-analysis of the perceptual boundary task, we find no evidence for an age-related decline in the slope. Within the recalibration tasks, the age effects are radically different. If proprioception declines with age, we would expect that the two recalibration tasks would yield very similar age-related effects because both tasks are based on reliability-based weighting of the sensory inputs (proprioceptive and visual). However, the integration task provided evidence for an age-related decline in proprioception, while, in the same population, the motor adaptation task did not. Note that the task-dependency extends to the somatosensory domain (large effects for touch vs limited ones for proprioception) and to the motor domain (large age-related effects for the Purdue Pegboard task but not for the visually-guided reaching movements).

The task-dependency extends further to an outcome-dependency. Indeed, in some tasks, one of the outcomes provided evidence for an age-related decline in proprioception (absolute error for the position matching task, bias in the perceptual boundary task) while another provided evidence against an effect of aging on proprioception (variability for the position matching task or slope for the perceptual boundary). The fact that one can choose what conclusion one makes about the presence of an effect of aging on proprioception or not makes it difficult to understand how a deficit in proprioception can differentially affect the different proprioceptive outcomes. If proprioceptive decline is as large as it is claimed, we should see a significant effect of age in all outcomes. To tackle this problem one needs to have a closer look at the outcomes and on the different factors that influence them.

In typical position sense tasks (e.g. arm position matching and perceptual boundary), proprioception can be operationalized as the systematic error between actual hand position and target, which we refer to as spatial accuracy or bias, or it can be operationalized as the inter-trial variability in actual hand location, which we refer to as consistency or sensitivity (Fig. 22). Our results show clear age-related effects in variables linked to bias (arm position matching – absolute error; perceptual boundary – bias), but those effects are less clear for variables related to consistency (arm position matching – variability; perceptual boundary – slope). This is highlighted by the results from the perceptual boundary task, in which we find a medium effect of age on bias, but a negligible effect on the slope (a measure of sensitivity). Yet, at this time, there is no consensus as to whether small bias (Fig.22.A) or small variability (Fig.22.B) represent better proprioceptive function.

**Figure 22.**
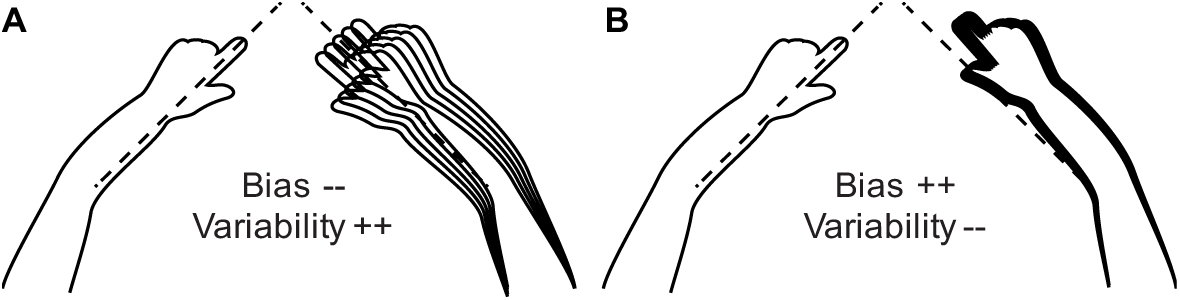
Representation of bias vs. variability. A) Low bias and high variability. B) High bias and low variability.

The interpretation of bias can be complicated because it can be affected by external factors during the experiment. In position sense tasks, proprioceptive judgements have to be made relative to the body midline (e.g., the arm position matching task and the perceptual boundary task). In many setups like ours, the location of the body midline is not precisely controlled. For example, the body midline can shift or rotate with respect to the workspace which would affect the bias but not the consistency (Fig. 22). Another reason to prefer consistency over bias is that, in typical sensory integration studies with proprioception, it is the variability and not the bias that is taken into account to weight the different sensory signals (van Beers et al., 1999). Finally, bias and consistency variables are differently impacted during high-level proprioceptive judgements (Héroux et al., 2024). High-level proprioception requires that spatial coordinates are transformed from one reference frame (body space) into an additional reference frame (visual space). This means that, for example in the case of visual targets, proprioceptive judgements require a reference frame transformation from the muscle space to the visual space. Importantly, Héroux and colleagues (2024) argue that in high-level proprioceptive judgements, large bias does not reflect impaired proprioception but rather differences in reference frame transformations. Consequently, the interpretation of bias in tasks with visual targets (as is the case in e.g. the perceptual boundary task) is complicated and not only related to the quality of proprioceptive function. To conclude, we cannot with certainty interpret the bias-related outcomes as measures of proprioceptive accuracy.

### Age-related differences are of limited size for typical position sense tasks

Looking back at our results while keeping the above remarks in mind, we find that, in our dataset, position sense tasks revealed limited age-related differences. This is in line with previous studies that found no age-related impairments in position sense (Batavia et al., 1999; Boisgontier & Nougier, 2013; Coffman et al., 2021; Djajadikarta et al., 2020; Dunn et al., 2015; Goble et al., 2012; Jordan, 1978; Pickard et al., 2003; Saenen et al., 2023; Schmidt et al., 2013; Wang et al., 2012; Weijs et al., 2024), and provides evidence against large age-related differences in position sense found by others (Adamo et al., 2007, 2009; Kaplan et al., 1985; Ko et al., 2015; Lord & Ward, 1994; Pai et al., 1997; Petrella et al., 1997; Skinner et al., 1984; Tulimieri & Semrau, 2023; Van Impe et al., 2012; Wingert et al., 2014; Yang et al., 2019). In some of the latter studies, small sample size probably leads to an overestimation of the effect of age on proprioception, consistent with the winner’s curse (Button et al., 2013). Studies with larger sample size have found significant age-related changes in position sense but these changes are of limited size (Herter et al., 2014). These are confirmed by our meta-analyses where we found that the effect of aging on position sense has a small effect size (Cohen’s d around d=0.25). If the effect size is so small, it means that most of the previous studies on the effect of aging on proprioception are vastly underpowered as most use 20 participants per age group, corresponding to a power of 10-15%.

However, our meta-analysis is only based on data from our laboratory. There are several possibilities about why position sense tasks in our laboratory could exhibit such limited effect size. One of them resides in the possibility that our population of older adults is more physically active or healthy than those recruited in other laboratories. Certainly they are more healthy than people from this age range in the general population (Harris et al., 2008; Martinson et al., 2010; van Heuvelen et al., 2005). It has been shown that physically active adults perform better in position sense tasks than inactive adults (Kitchen & Miall, 2019, 2021). Indeed, higher levels of physical activity can prevent a decline in proprioception with age (Ribeiro & Oliveira, 2007). Since we don’t have data on the level of physical fitness of our participants, we cannot exclude the possibility that group differences can be explained by the level of physical fitness. Future studies with measurement of the level of physical fitness are recommended. Yet, this explanation would not account for the task-dependency that we observe.

### Task-dependency of age-related effects might be explained by cognition

Some tasks seem to consistently show an effect of age, such as the haptic perception task (Overvliet et al., 2013; Saenen et al., 2023; Wang et al., 2012). One could thus conclude that haptic perception declines with age. Interpretation of such age-related differences, however, should be done with care because of the higher level of complexity in the stimuli, instructions and execution of this task. The haptic proprioception task is a high-level sensory processing task, involving more than pure proprioception. As such, several aspects of cognition could influence performance on this task. Indeed, proprioceptive encoding is more likely to be compromised in older adults with low working memory capacity (Goble et al., 2012). For instance, working memory, which is lower in our older participants, plays an important role in this task as the shape has to be memorized and mirrored before being reproduced with the contralateral arm. While taking into account individual differences in visuo-spatial working memory capacity does not suppress the age effect, age-related changes in other cognitive functions might account for this effect. This can possibly be explained by the interference between visuo-spatial and proprioceptive memory capacity when concurrently performing a proprioceptive and a spatial task (i.e., in the case of the haptic perception task, mirroring the shape to the contralateral side) (Horváth et al., 2022). While proprioception does certainly play a role in this task, it is unknown how much the age-related decline in performance is due to a decline in proprioception.

Age-related differences in cognition might also account for the limited effect size of aging in the position matching task. Indeed, dual-task paradigms have provided evidence that cognitive interference affects proprioceptive consistency and processing speed (Bigongiari et al., 2018; Jiang et al., 2023). One dual-task study found an improvement when proprioceptive matching in a single joint was tested in combination with a cognitive dual-task, but a worsening when whole arm matching was combined with the cognitive dual-task (Ager et al., 2024). Taken together, these findings suggest that cognition influences performance in proprioceptive tasks, certainly when task complexity increases. Consequently, we need to carefully consider the possible concurrent effects of cognition when interpreting age-related effects in proprioceptive tasks. Even more so, cognition might explain some of the task-dependency of aging effects that we find in this study.

### The modality of the answer might explain task-dependency

Some of the task-dependency can arise from the fact that the modality of report differs in nature. While many of our tasks are perceptual tasks based on conscious verbal report (e.g. position sense and perceptual boundary tasks), other tasks from our task battery are motor tasks where proprioceptive information is used to shape the motor output (e.g. integration task). In the latter, conscious reporting is eliminated, thereby reducing strategy use and cognitive demand. This distinction is similar to the idea of Milner & Goodale (2008) who found that visual information is processed differently for perception and action. This difference in processing has been illustrated by illusion studies where an objects illusory size or weight does not influence the subjects grip aperture or force, but it does influence perceptual judgements about its size or weight (Aglioti et al., 1995; Flanagan & Beltzner, 2000; van Doorn et al., 2007). The distinction has also been made for tactile processing (Pruszynski et al., 2018). Performance in the motor tasks (i.e. implicit adaptation and integration) is correlated, suggesting that proprioception is used in a similar way in both tasks.

Larger age-related effects in the recalibration tasks (integration – slope; implicit adaptation – result of meta-analysis), reveal that perturbed visual feedback has a larger effect on behavior for the older adults than for the young adults. This is consistent with previous findings in similar adaptation tasks (Cisneros et al., 2024; Vachon et al., 2020), in shape recognition where older adults are more sensitive to disturbed visual feedback during limb movement (Wang et al., 2012), and in virtual reality where older adults are more likely to experience a virtual hand as their own (Risso et al., 2024). These findings suggest that older adults will show higher levels of recalibration following discrepancies between proprioceptive and visual information.

According to Bayes optimal integration rule, proprioceptive and visual information are weighted depending on their reliability (Ernst & Banks, 2002; van Beers et al., 1999). Thus, the higher bias towards vision in our group of older adults should entail that a lower weight is being given to proprioceptive information because of its lower consistency, as was indeed found by Risso and colleagues (2024). Two of our findings, however, are difficult to reconcile with this framework of reliability-based weighting of sensory information and reinforce our idea that there exist two separate streams for proprioception. The effect of age on the reliability/consistency of proprioception as measured in tasks involving conscious reports is limited (position matching task and perceptual boundary task). Therefore, we would expect a limited effect of age on the weighting of proprioceptive and visual information. Yet, the integration task, which is a proprioception-for-action task, did exhibit large age-related differences. The absence of a link between the reliability of proprioception measured with conscious reports and the weighting of vision and proprioception could stem from the existence of two separate streams for action and perception. But there are alternative explanations.

One possible explanation is that the weight given to sensory information is unrelated to its reliability. Indeed, previous findings pose that attenuated proprioception alone cannot explain the up weighting of visual information (Kitchen & Miall, 2021; Parthasharathy et al., 2022; Vachon et al., 2020; Vandevoorde & Orban de Xivry, 2021). Mikula and colleagues (2018) found that weighting of sensory inputs is not reliability-based, but rather learned and modality-specific. Even more, Block & Bastian (2011) pose that integration of visual and proprioceptive inputs can be driven by reliability-based weighting, but that the realignment of both inputs can be flexibly adapted according to the circumstances such as the brevity of the misalignment or attentional resources. Alternatively, the higher weighting of visual information could be explained by the prior belief that the visual and proprioceptive inputs represent the same object, namely the hand. Following the Bayesian Causal Inference model as described by Risso (2024), this prior belief is a top-down cognitive component and is independent of reliability of proprioception. Notably, when the prior believe that hand and cursor share a common cause is stronger, the sensory inputs could each be assigned a higher level of redundancy. In previous work by Laurienti and colleagues (2006) older adults showed enhanced integration of redundant information which could explain relative up weighting of visual information by older adults.

### The different outcomes do not reflect a single construct

In addition to the fact that results within the same domain (sensory/recalibration/motor) do not provide clarity about age-related differences, we find that relationships between parameters that should measure the same construct are negligible or, at best, small. Concerning proprioception, we only find an association between consistency in arm position matching and perceptual boundary. For all other proprioceptive variables, we found weak and non-significant correlations. This finding supports the idea that consistency measures are preferable over bias measures. Concerning recalibration, we found that the outcomes from both tasks did correlate even though the effect of age is radically different in both tasks.

Any correlations we didn’t detect in our sample because of a lack of power will probably not be substantial. It is assumed that all these tasks measure some aspects of proprioception (the latent factor), but either these tasks measure different aspects of proprioception (low-level vs. high-level, perception vs. action, etc.) or proprioception contributes only minimally to task performance. The lack of association between proprioceptive variables is in accordance with previous studies showing no associations between outcome variables from different proprioceptive tasks (Horváth et al., 2023; Lowrey et al., 2020; Nagai et al., 2016). Also, Horváth and colleagues found that consistency and bias are not associated within the elbow and the knee (2024). Interestingly, outcomes from the position matching task that exhibited similar age-related effects (d=0.25 and 0.24) were also not correlated.

## Conclusion

Our battery of proprioception-related tasks suggests that age-related changes in proprioception are not as clear as considered. They are task-dependent, outcome-dependent and are of limited size when they are actually present. In addition, some of the reported age-related effects on proprioception are probably confounded by other factors such as cognition, reference frame transformation and perceptual biases. Our dataset suggests that consistency metrics should be preferred to accuracy ones. There is a need to understand the lack of associations between outcome variables from standard proprioceptive tasks. Our data and that of others (Horváth et al., 2023, 2024; Lowrey et al., 2020; Nagai et al., 2016) suggests that we can’t get a good measure of proprioception via the current standard tasks. Such measure of the construct of proprioception is required to make valid conclusions about age-related effects.

## DATA AVAILABILITY

All data used for statistical analysis can be found on the RDR repository of the KU Leuven: https://rdr.kuleuven.be/. Raw kinematic data for each task can be obtained from the last author upon reasonable request.

All analysis scripts can be found at: https://rdr.kuleuven.be/.

## GRANTS

This work was supported by the FWO grant G001724N.

## DISCLAIMER

The funder had no role in study design, data collection and analysis, decision to publish, or preparation of the manuscript.

## DISCLOSURE

No conflicts of interest, financial or otherwise, are declared by the authors.

## AUTHOR CONTRIBUTION

S.V.D.P., and J-J.O.d.X. conceived and designed research; S.V.D.P performed experiments; S.V.D.P and J-J.O.d.X. analyzed data; S.V.D.P, and J-J.O.d.X. interpreted results of experiments; S.V.D.P and J-J.O.d.X. prepared figures; S.V.D.P drafted manuscript; S.V.D.P edited and revised manuscript; S.V.D.P, and J-J.O.d.X. approved final version of manuscript.

## ACKNOWLEDGMENT

We thank Lize Geudens and Gwendoline Nore for help with data collection.

## Notes

https://rdr.kuleuven.be/

### Competing Interest Statement

The authors have declared no competing interest.

### Summary of Updates

Superiority index added to results section; Discussion updated to make the overall messages more clear.

