## Supplemental Table 1 for "Age-related changes in proprioception are of limited size, outcome-dependent and task-dependent"

### Supplementary material

*Table 1.* Factor analysis on the six position sense outcome variables.

| Variable | Factor loading |
|---|---|
| Arm position matching accuracy | -0.09 |
| Arm position matching variability | -0.41 |
| Perceptual boundary bias | 0.31 |
| Perceptual boundary sensitivity | 0.43 |
| Shape reproduction accuracy | 0.38 |
| Integration consistency of return (0°) | -0.09 |
